# Periodic spatial patterning of gene expression via a natural quorum-synchronized oscillator

**DOI:** 10.64898/2026.11.13.766566

**Authors:** Varun Sridhar, Steven Guenther, Kyle Guo, Yein Ra, Kilian Dekoninck, Marcos M. Vasconcelos, Matthew F. Traxler

## Abstract

In many biological systems, genetic oscillators that function in growing tissues can establish spatially repetitive patterns of gene expression. However, few mechanisms for generating such patterns are known in bacteria. Here, we describe an oscillator within the quorum sensing circuit of *Streptomyces coelicolor*. Through modeling and genetic experiments, we demonstrate that this quorum oscillator controls periodic expression of a gene involved in antibiotic biosynthesis at the periphery of expanding colonies. We show that environmental factors, namely small molecule diffusabilty and carrying capacity, independently modulate the oscillator frequency and amplitude, respectively. The architecture of the quorum oscillator imbues this ‘clock’ with three emergent features; it is synchronized locally across cells, only advances due to collective cellular communication, and its dynamics are directly responsive to environmental conditions.

## Introduction

Biological oscillators enable a wide range of rhythmic behaviors in organisms across the tree of life (1–5). When coupled to tissue growth, oscillatory gene expression can generate periodic spatial patterns that underlie the formation of repetitive biological structures (6–8). Some key eukaryotic examples include vertebrate somitogenesis and arthropod segmentation (9–11), in which oscillating genetic circuits direct the sequential formation of body segments, and the *Arabidopsis* root clock, which periodically establishes sites of lateral root formation (12, 13). In the most well-characterized of these systems, vertebrate somitogenesis, intercellular signalling plays an important role in coordinating the genetic oscillations across cells to generate the foundational periodic spatial patterns (9). How bacteria may employ analogous genetic oscillators to build periodic spatial patterns remains much less explored.

Several natural oscillatory systems have been described in bacteria. These include circadian cycles in cyanobacteria via the post-translational KaiABC oscillator (14, 15) and cross-generational fluctuations of cyclic di-GMP concentrations in *Pseudomonas aeruginosa*, which arise via an unknown mechanism (16). Additionally, nitrogen stress response cycles in *Bacillus subtilis* biofilms have been shown to trigger cell-autonomous oscillations that can lead to periodic spatial segmentation (17). While these behaviors involve oscillating components, and in the case of *B. subtilis* can lead to periodic spatial patterns, none of these systems rely on intercellular communication to synchronize oscillatory functions across cells.

The past three decades have seen the deployment of a wide variety of synthetic genetic oscillators in bacteria. Key among these is an archetypal circuit known as the repressilator, in which three transcriptional regulators are arranged in a cyclic repression network that can generate sustained oscillations (18). However, repressilator activity proceeds asynchronously across cells and is often highly variable in terms of frequency and amplitude (18–21). Quorum sensing is a well-known mechanism that can coordinate changes in gene expression across populations of bacterial cells (22), and the synchronicity of synthetic systems has been improved by incorporating quorum control at different stages within oscillating circuits (23–30). These synthetic systems and a substantial body of theoretical evidence (23–26) demonstrate that synchronized genetic oscillators can be built in bacteria, but natural examples of such circuits have yet to be identified.

Here we report that the bacterium *Streptomyces coelicolor* employs a natural genetic oscillator to produce periodic, concentric patterns of gene expression within growing colonies. As colonies expand, this circuit generates recurrent pulses of expression of a key antibiotic-biosynthesis regulator at the advancing colony periphery, leaving periodic rings of expression in its wake. The circuit that enables this behavior contains a cyclic repression network reminiscent of synthetic repressilators, but also incorporates a quorum-sensing gate within the oscillator itself. Progression through this quorum-gated transition requires accumulation of the diffusible intercellular signal, thereby synchronizing the oscillator across the collective, and imbuing the circuit with the ability to modulate frequency and amplitude in response to environmental conditions. Our findings reveal a naturally evolved strategy in which intercellular communication synchronizes genetic oscillations to generate periodic spatial patterns in a growing bacterial community.

## Results

### The *S. coelicolor* QS circuit

While the disparate connections between each component of the *S. coelicolor* QS circuit have been previously elucidated, a holistic framework with easily identifiable regulatory motifs has yet to be assembled (31). Here, we present such a network (Fig. 1A). QS in *S. coelicolor* is mediated by γ-butyrolactone (GBL) signal molecules that are made by the synthase ScbA (32). These signals are presumed to diffuse freely in and out of cells (33). Upon accumulation to a threshold concentration, GBLs bind to and inhibit the DNA-binding activity of the cognate receptor-regulator, ScbR (32). In its ligand-unbound state, ScbR actively binds DNA and represses the transcription of many genes including itself. A key target of ScbR repression is *scbR2*, which encodes a related receptor-regulator protein that does not respond to GBL signals (34, 35). ScbR2 represses *scbA* transcription, closing a negative feedback loop (36). This negative feedback loop consists of ScbA producing GBLs, GBLs inhibiting ScbR (upon their accumulation to a threshold concentration), ScbR repressing the transcription of *scbR2*, and ScbR2 in turn repressing the transcription of *scbA*. The *cpkO* gene is a downstream target of this negative feedback loop, since its transcription is repressed by either ScbR or ScbR2 (37). CpkO is a pathway-specific regulator that activates transcription at multiple promoters within the *cpk* gene cluster, leading to production of coelimycin antibiotics (31, 38, 39).

**Fig. 1.**
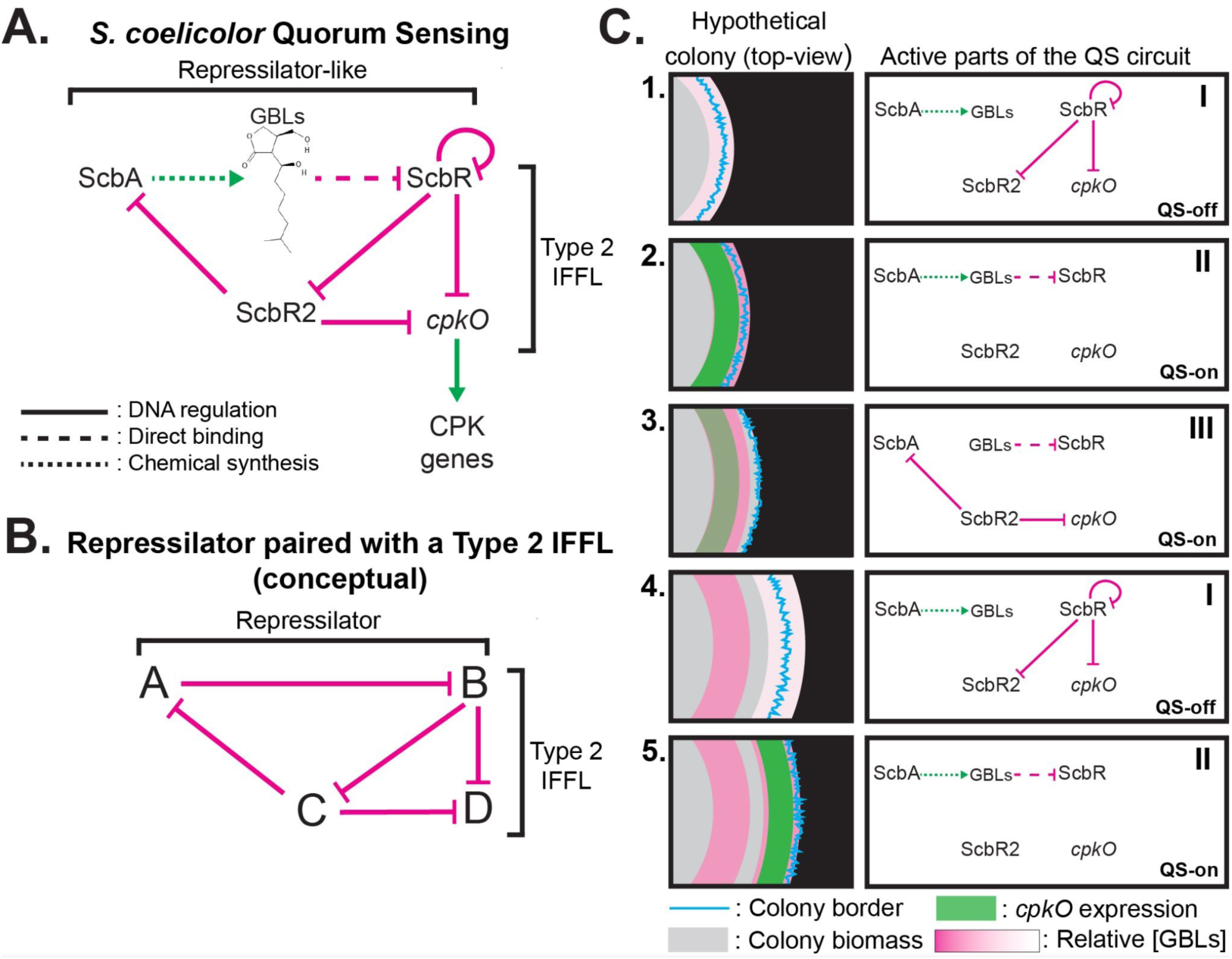
The *S. coelicolor* quorum oscillator. **(A)** A network diagram of the *S. coelicolor* quorum sensing circuit, dubbed ‘quorum oscillator’. **(B)** Conceptual mockup of a repressilator circuit integrated with a Type 2 IFFL. Solid arrows represent activity through DNA regulation, dashed arrows represent activity through direct binding to a target, and dotted arrows indicate direct manipulation of molecules. Green arrows indicate a positive control mechanism (upregulation/activation/synthesis), and magenta flat arrows indicate a negative control mechanism (repression/inactivation). **(C)** Conceptualization of how the QS circuit may manifest on a spatial axis as colonies grow. This is a mockup of a top-down view of a colony growing. We show three distinct steps of the QS circuit and the growing colony edge (blue line) at each circuit, alongside conceptual relative locations of GBL concentration (magenta area) and *cpkO* expression (green area). We hypothesize that these three steps occur repeatedly at the growing colony edge. Each stage of the QS circuit shown is a hypothesis of what components are active at the colony edge in each stage.

The topology of the QS network can be understood as a quorum-gated repressilator-like circuit consisting of ScbA, ScbR, and ScbR2 (Fig. 1B) paired with a Type 2 incoherent feed forward loop (IFFL) (40) consisting of ScbR and ScbR2 controlling *cpkO* expression. An IFFL is a well-known regulatory motif whose activity results in a singular transient pulse of gene expression. The coupling of a pulse-generating IFFL with an oscillator in this way suggests that a pulse of *cpkO* expression will occur with each oscillation. This topology prompts the hypothesis that a pulse of *cpkO* expression will occur after GBLs accumulate to a threshold concentration, but before ScbR2 has accumulated to a high enough level to repress expression of *cpkO*.

We next considered a conceptual model of how this circuit might function to pattern gene expression in growing colonies of *S. coelicolor* (Fig. 1C). When a colony is small and at a low population density, the cells are in a QS-off state (Fig. 1C, Panel 1). As the colony grows, it accumulates GBLs and eventually reaches a QS-on state, generating a pulse of *cpkO* expression (Fig. 1C, Panel 2). In the QS-on state, transcriptional repression of *scbR2* is relieved, and accumulating ScbR2 protein begins to repress *scbA* transcription (Fig. 1C, Panel 3) turning off GBL production. Although their production has been turned off, the GBLs themselves remain present in the extracellular environment; therefore, the cells remain in a QS-on state. As the colony grows, cells at the periphery eventually expand into new space that lacks GBLs (Fig. 1C, Panel 4). In the absence of GBLS, newly made ScbR once again represses its own transcription and transcription of *scbR2*, resetting the oscillator to its initial state. We thus hypothesized that the architecture of the *S. coelicolor* QS circuit would result in repetitive oscillations of *cpkO* expression localized to the growing colony edge at regularly spaced intervals. We refer to this circuit as the *S. coelicolor* ‘quorum oscillator’.

### Periodic spatial patterning of a quorum-controlled gene at the *S. coelicolor* colony edge

Based on the topology of the *S. coelicolor* quorum oscillator, we hypothesized that *S. coelicolor* colonies could generate periodic pulses of *cpkO* gene expression at their growing edge. To test this prediction, we sought to visualize *cpkO* expression in *S. coelicolor* colonies. We did so by monitoring the promoter activity of *cpkO* with an EGFP-based transcriptional reporter fusion (P*_cpkO_-egfp*) integrated into the *S. coelicolor* genome in single copy (41). We also inserted a construct that constitutively expresses the red fluorescent protein mCherry under control of the strong promoter P*_ermEp*_* (P*_ermEp*_-mCherry*) to enable normalization for colony growth (GFP/RFP) (42). WT colonies expressing these two fluorescence constructs were grown for 42 hours to a size of ∼500 microns in diameter before being monitored for 54 hours with time-lapse fluorescence microscopy.

Consistent with our hypothesis, we observed periodic pulses of *cpkO* expression in colonies at their periphery (Fig. 2A, Movie S1). To visualize this activity over time, we generated a kymograph that tracks normalized *cpkO* expression along the radius of this colony throughout the time course (Fig. 2B). The pulses of normalized *cpkO* expression are seen in this kymograph as red zones at the colony edge, highlighted by the dashed boxes. This pattern was consistent across three replicates (Fig. S10). To further quantify this pulsing behavior, we measured normalized *cpkO* expression in a defined band at the growing colony edge, in triplicate. Plotting the phase-aligned averaged expression over time revealed four clear oscillations during the observation interval (Fig. 2C), with an average period of 12.21 hours and average amplitude of 0.13 normalized fluorescence intensity units. In contrast, the promoter for a different gene, *bldN*, which encodes a sigma factor that regulates morphological development in *S. coelicolor* and is not controlled by QS, did not exhibit oscillatory expression (Fig. S2). Additionally, we found that expression of *scbA*, the GBL synthase, was very low in this experiment setup in comparison to *cpkO* (Figure S3). These results are broadly consistent with the hypothesis that the QS network of *S. coelicolor* can structure periodic spatial patterns of *cpkO* expression at the colony periphery.

**Fig. 2.**
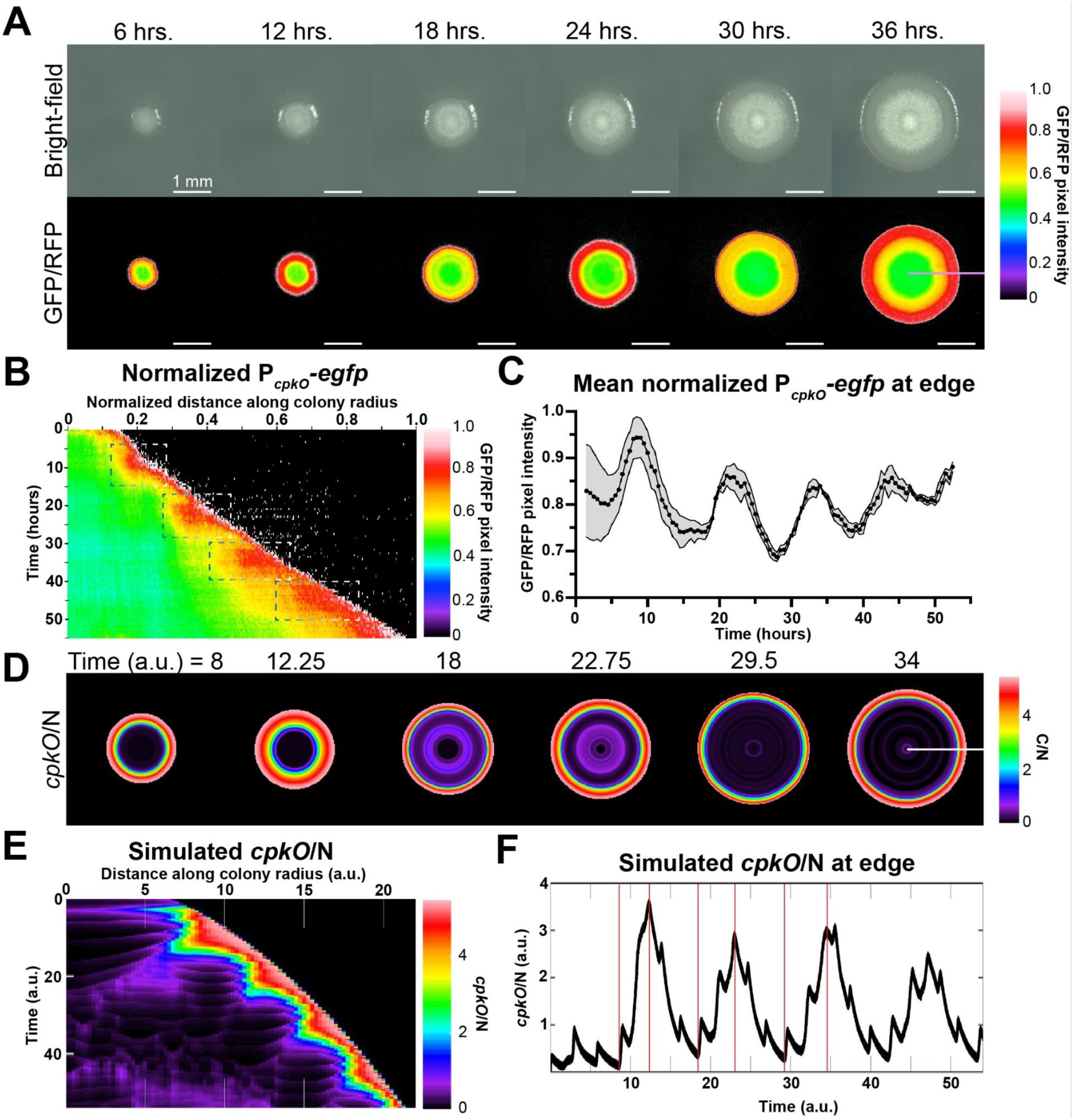
*cpkO* expression pulses cyclically at the edge of simulated and real *S. coelicolor* colonies. **(A)** A series of images showing a representative WT colony growing (BF, top) and expressing a ratio of P*_cpkO_-egfp*/P*_ermEp*_-mCherry* (GFP/RFP, bottom) over 36 hours. Images were curated from a representative time-lapse movie. Time points were selected based on local minimums and maximums of the observed oscillation. The color bar represents the LUT on which the GFP/RFP images are projected from. The mauve line in the final GFP/RFP image indicates the radius that the kymograph in (B) was built on. Scale bars = 1 mm **(B)** A kymograph that charts P*_cpkO_-egfp*/P*_ermEp*_-mCherry* expression along the radius of the simulated colony (horizontal axis) over time (vertical axis). Each of the four pulses of normalized *cpkO* expression is highlighted by the dashed boxes. **(C)** Normalized fluorescence intensity plot of mean P*_cpkO_-egfp*/P*_ermEp*_-mCherry* expression at the edge of colonies over time. Data points are an average of three colonies, and shading represents SE. SE for some points may not be visible. **(D)** A series of images showing simulated *cpkO*/N expression in a virtual colony, based on a mathematical model of our proposed QS circuit. Time *(figure caption, continued)* points were selected based on local minimums and maximums of the observed oscillation. The white line in the final panel represents the radius that the kymograph in (E) was built on. **(E)** A kymograph that charts *cpkO*/N expression along the radius of the simulated colony (horizontal axis) over time (vertical axis). **(F)** A plot of mean simulated *cpkO*/N in a narrow band at the growing colony edge. Red lines indicate the time points at which images in (D) were selected.

### Mathematical modeling of the quorum oscillator predicts periodic spatial patterning of *cpkO* expression at the colony edge

To see if a circuit with the topology of the *S. coelicolor* quorum oscillator is sufficient to enable periodic spatial oscillations of *cpkO*, we designed a mathematical model of the circuit in Fig. 1A. This model integrates six partial differential equations (PDEs) to simulate spatial and temporal *cpkO* expression in a *S. coelicolor* colony expanding in a virtual 2D space. These PDEs are summarized in Fig. S1, and detailed in the Materials & Methods. Model parameters were tuned using a hybrid approach based on both reported regulatory relationships (43, 44) and data generated here.

The model predicted *cpkO* expression (normalized to number of cells, N) in a periodic spatial pattern at the virtual colony edge (Fig. 2D, Movie S2). A kymograph of the simulation output enables clear visualization of the *cpkO*/N pulses at the virtual colony edge over a set time interval (Fig. 2E). Measuring *cpkO*/N expression in a defined band at the virtual colony edge further demonstrates this oscillating behavior (Fig. 2F). We note that the model also predicted a set of minor oscillations superimposed on the major oscillatory activity of the circuit. Due to limits in resolution in both time and fluorescence sensitivity, our experimental data do not address these minor oscillations. Thus, we limit our interpretation to the activity captured in the major oscillations predicted by our mathematical model. Taken together, these simulation results demonstrate the sufficiency of the *S. coelicolor* QS circuit to generate oscillatory patterns of gene expression that broadly align with those observed in living colonies.

### Strains lacking ScbA or ScbR fail to produce periodic spatial expression patterns

We next sought to test the hypothesis that the *S. coelicolor* quorum oscillator is the driver of the observed cycles in *cpkO* expression. To do so, we checked *cpkO* expression in Δ*scbA* and Δ*scbR* cells both in simulations and *in vivo*, predicting that *cpkO* transcription would not oscillate in strains lacking these key QS components. In *in silico* knockout mutants (i.e. simulations where the production rate of either ScbA or ScbR (α_A_ or α_R_, respectively) was set to zero), normalized *cpkO* expression was observed at the colony edge, and did not exhibit oscillations. This can be seen in kymographs that chart *cpkO*/N expression along the radius of the colony (Figs. 3A, 3D). These simulations predicted a comparatively thinner band of expression at the periphery of colonies lacking ScbR.

**Fig. 3.**
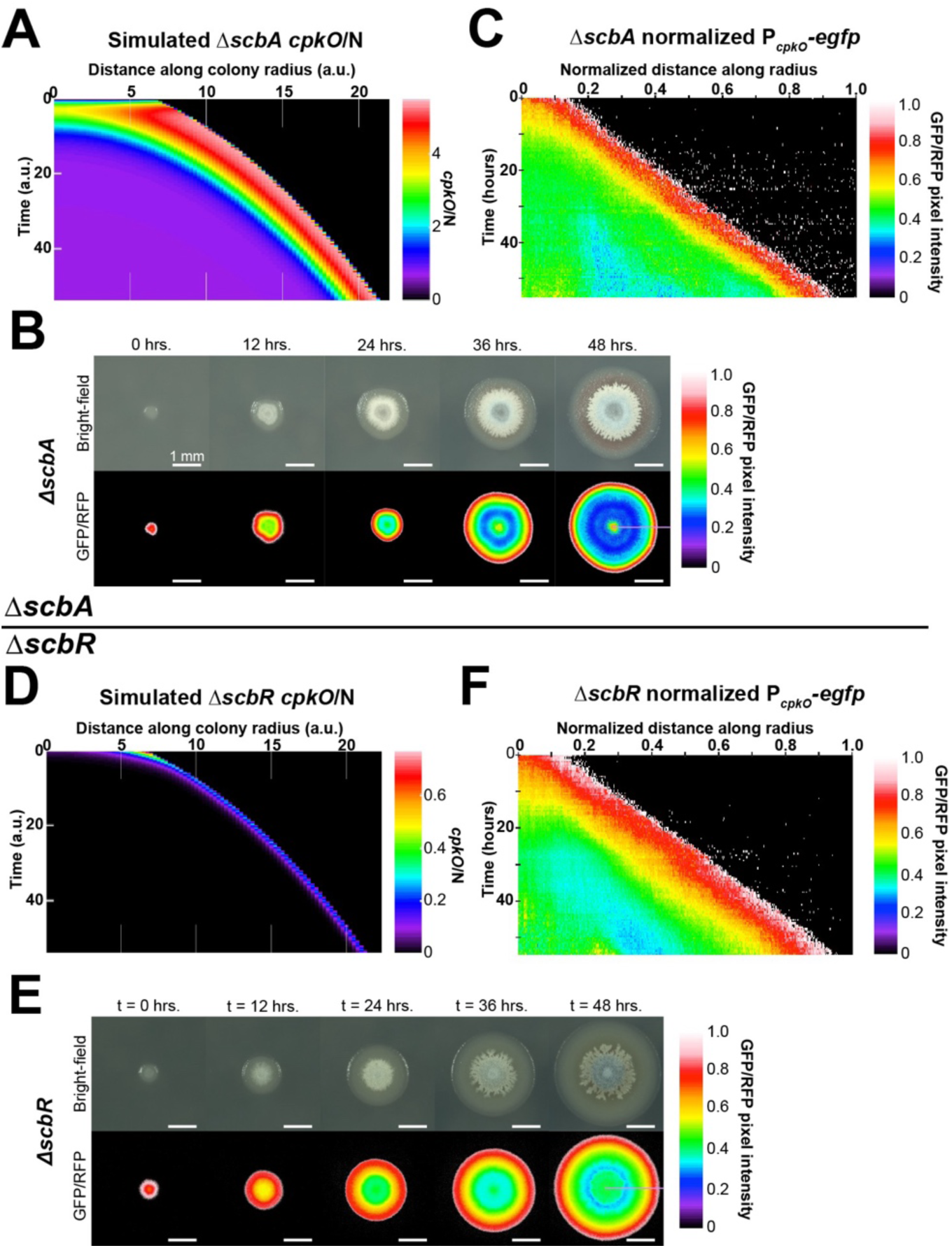
Pulses of normalized *cpkO* expression are abolished in locked “QS-off” mutants. (A, **D)** Kymographs charting *cpkO*/N expression along the radius of a simulated Δ*scbA* (A) or Δ*scbR* (FD colony. **(B, E)** Representative image time series of a real Δ*scbA* (B) or Δ*scbR* (E) colony growing over time. Images were curated from representative time-lapse movies. BF (top) and GFP/RFP (bottom) channels are shown. The color bar represents the LUT on which the GFP/RFP images are projected from. The mauve line indicates the axis on which the kymographs in C and F were built. Scale bars = 1 mm. **(C, F)** Kymographs that chart P*_cpkO_-egfp*/P*_ermEp*_-mCherry* (GFP/RFP) expression along the radius of a representative Δ*scbA* (C) or Δ*scbR* (F) colony.

We next made *in vivo* knockout strains lacking *scbA* or *scbR* and monitored P*_cpkO_-egfp*/P*_ermEp*_-mCherry* expression. Similar to simulated colonies, normalized *cpkO* expression in both mutant strains backgrounds was seen at the colony periphery, and did not exhibit oscillatory expression profiles (Fig. 3B and C, and 3E and F, and S4B and D). We noted that in the living Δ*scbA* and Δ*scbR* strains, the width of the band of sustained normalized *cpkO* expression at the colony periphery was roughly the same. In contrast, the band of *cpkO* expression in the simulated Δ*scbR* colony was thinner. These simulated and *in vivo* results indicate that, at a qualitative level, normal functions of ScbA and ScbR are both required for the oscillatory expression of *cpkO* at the colony periphery in the WT.

We next considered if the sustained peripheral expression of *cpkO* in the *scbA* and *scbR* mutant strains might arise from the relative repressive strengths of ScbR and ScbR2 on their respective targets within the QS circuit. To explore this possibility, we tested many values of the half-saturation constants associated with ScbR or ScbR2’s repressive strengths (K_RR_. K_RR2_, K_RC_, and K_R2C_). We found that in the simulated *scbA* mutant, the thickness of the band of *cpkO* expression was strongly influenced by the repressive strength of ScbR on itself (K_RR_), and on ScbR2 (K_RR2_) (Fig. S5A, B). Furthermore, reducing the simulated repressive strength of ScbR on *cpkO* (K_RC_) led to strong *cpkO* expression across the entire colony (Fig. S5C). In the simulated Δ*scbR* colony, the very thin band of normalized *cpkO* expression at the periphery (Fig. 3D) also became wider when we lowered the repressive strength of ScbR2 on *cpkO* expression (K_R2C_) (Fig. S6). These results are consistent with the idea that, in a circuit lacking the QS signal or receptor, normalized expression of the downstream target (i.e. *cpkO*) is relegated to a peripheral sustained band whose dimensions are dictated by the relative repressive strengths of ScbR and ScbR2 at specific promoters within the QS circuit.

### Strains lacking ScbR2 exhibit a single peripheral pulse followed by sustained expression of *cpkO*

We next considered the role of ScbR2 in the function of the *S. coelicolor* quorum oscillator (Fig. 1A). We hypothesized that Δ*scbR2* colonies would advance normally through the QS circuit to the point of expression of *cpkO*. However, we anticipated that in Δ*scbR2* colonies, *scbA* expression would not be subsequently repressed and the production of GBLs could continue unimpeded. To explore the implications of this altered regulation, we checked the pattern of normalized *cpkO* expression in a simulated *scbR2* mutant colony (i.e. where the production rate of ScbR2 (α_R2_) was set to zero. This simulation showed a rapid pulse of normalized *cpkO* expression at the colony edge followed by a pronounced decrease in expression. After this transient attenuation, *cpkO* expression returned to an elevated, sustained level at the colony periphery (Fig. 4A, B). Next, we knocked out *scbR2* and monitored normalized *cpkO* expression (Fig. 4C) *in vivo*. Similar to the model, the *scbR2* mutant exhibited a single pulse of normalized *cpkO* expression followed by sustained, elevated expression at the colony periphery (Fig. 4C-E). While the relative magnitudes of the initial pulse of *cpkO* expression varied between the model and the living colony, the qualitative pattern was clear in both cases.

**Fig. 4.**
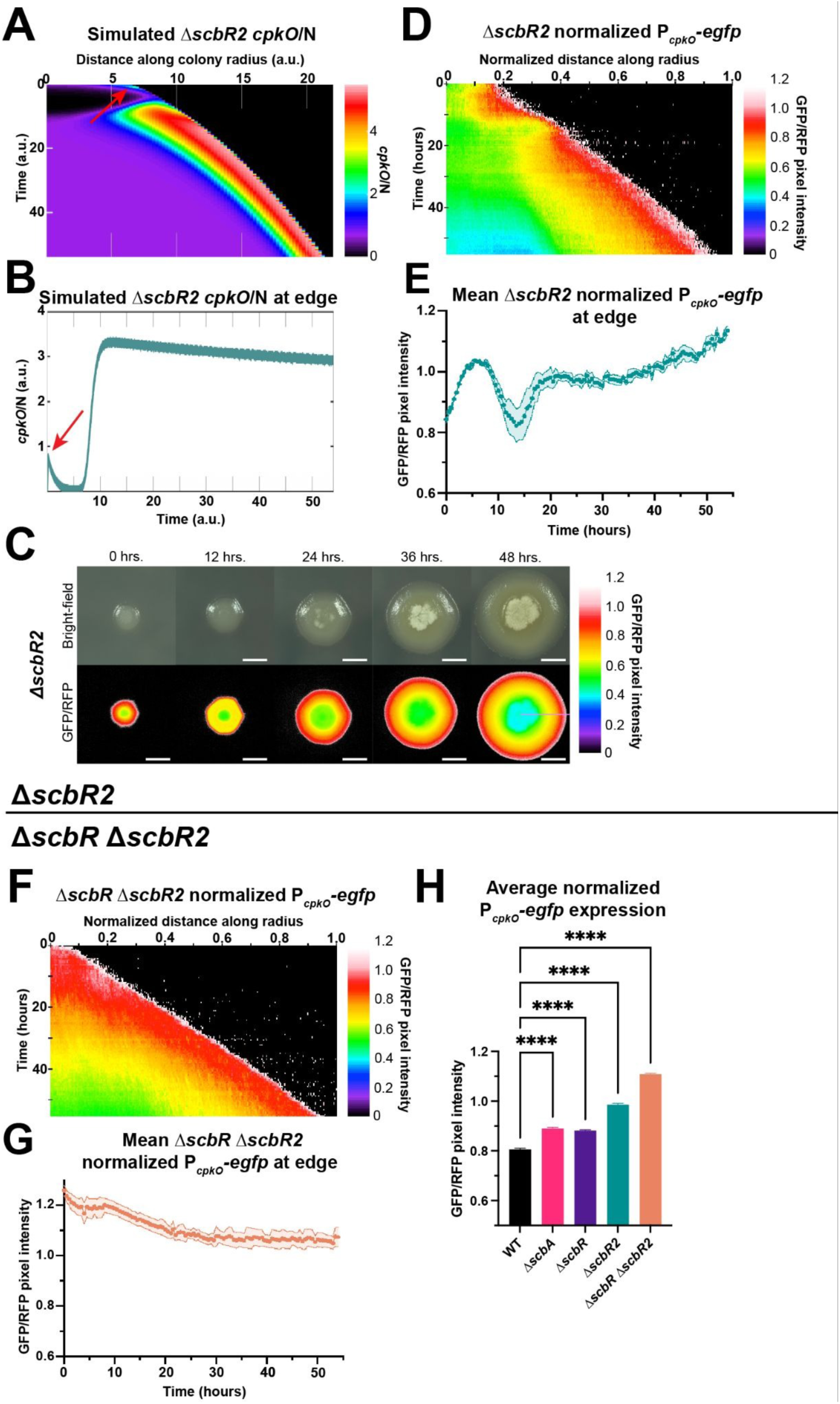
Pulses of normalized *cpkO* expression are largely disrupted in a locked “QS-on” mutant. **(A)** Kymograph charting *cpkO*/N expression along the radius of a simulated Δ*scbR2* colony. **(B)** Measured *cpkO*/N in a narrow band at the simulated Δ*scbR2* colony edge. The arrow indicates the observed spike of simulated *cpkO*/N expression. **(C)** Representative image time series of a real Δ*scbR2* colony growing over time. Images were curated from representative time-lapse movies. BF (top) and GFP/RFP (bottom) channels are shown. The color bar represents the LUT on which the GFP/RFP images are projected from. The mauve line indicates the axis on which the kymograph in D was built. Scale bars = 1 mm. **(D, F)** Kymographs that chart P*_cpkO_-egfp*/P*_ermEp*_-mCherry* (GFP/RFP) expression along the radius of a representative Δ*scbR2* (D) or Δ*scbRΔscbR2* (F) colony. **(E, G)** Fluorescence intensity plot that graphs GFP/RFP in a narrow band at the edge of Δ*scbR2* (E) or Δ*scbRΔscbR2* (G) colonies. The line denotes the mean of three measured colonies, with shading representing SE. SE for some data points may not be visible. **(H)** Average normalized P*_cpkO_-egfp* expression across an entire time course, in triplicate, for every measured strain. Significant differences between means compared between indicated strains were determined using an ANOVA. “****” indicates p ≤ 0.0005.

We next considered what might explain the singular pulse of *cpkO* expression in Δ*scbR2* colonies. As in the first cycle in the WT, GBLs deactivate the repression mediated by ScbR when they accumulate. This likely leads to increased expression of both *scbR* itself and *cpkO*. However, without ScbR2 to subsequently repress production of *scbA*, production of both GBLs and ScbR may continue unabated. We hypothesize that the strong production of ScbR might lead to a single period in which ScbR accumulation outpaced GBL production, leading to a transient ScbR-mediated repression of *cpkO* expression. This pause in expression might then be relieved after some time, during which GBL accumulation might reach a point that enabled a sustained inactivation of ScbR, leading to continuous *cpkO* expression at the colony periphery.

If the pause in *cpkO* expression did result from the transient activity of ScbR following its accumulation, then a clear prediction is that this transient attenuation would not occur in a mutant lacking both ScbR and ScbR2. Furthermore, we expected that a Δ*scbR*Δ*scbR2* strain would exhibit abnormally high expression of *cpkO* in the absence of both repressors. To test these predictions, we monitored P*_cpkO_-egfp*/P*_ermEp*_-mCherry* expression in a Δ*scbR*Δ*scbR2* strain. As expected, the Δ*scbR*Δ*scbR2* strain did not exhibit an initial transient pulse of normalized *cpkO* expression, rather it showed sustained *cpkO* expression at the periphery of the colony (Fig. 4F, G). Notably, the Δ*scbR*Δ*scbR2* strain had the highest average normalized *cpkO* expression at the colony periphery of all strains tested (Fig. 4H). These results are consistent with the idea that ScbR2 functions to reset the QS circuit to its initial “off” state at the expanding colony periphery. Additionally, while ScbR and ScbR2 both play a role in the repression of *cpkO*, the order of their activity within the circuit (i.e. ScbR active first, and ScbR2 active subsequently) leads to different effects on *cpkO* expression in strains lacking these regulators.

### Predictive simulations indicate that GBL diffusabilty and carrying capacity independently modulate the periodicity or amplitude of *cpkO* oscillations

The experiments above have demonstrated the role of specific genes in the function of the *S. coelicolor* quorum oscillator. We next sought to determine how environmental constraints might influence attributes (namely the periodicity and amplitude) of the oscillations of *cpkO* expression. Here we focused on two parameters determined by the local environment: D_G_, or the diffusion of GBLs (as a function of the density of the environment), and K_N_, or carrying capacity (as a function of nutrient availability).

To probe these two parameters, we ran a series of 20 simulations with a range of D_G_ or K_N_ values and assessed the effect on the resulting average periodicity and amplitude values (Fig. 5). The simulations indicated that increasing the GBL diffusabilty (D_G_) led to increased periodicity, while decreasing the carrying capacity (K_N_), and hence the density of cells in the simulated colony, led to a decrease in the amplitude. These model outputs suggest that environmental factors could independently influence key aspects of oscillating *cpkO* expression.

**Fig. 5.**
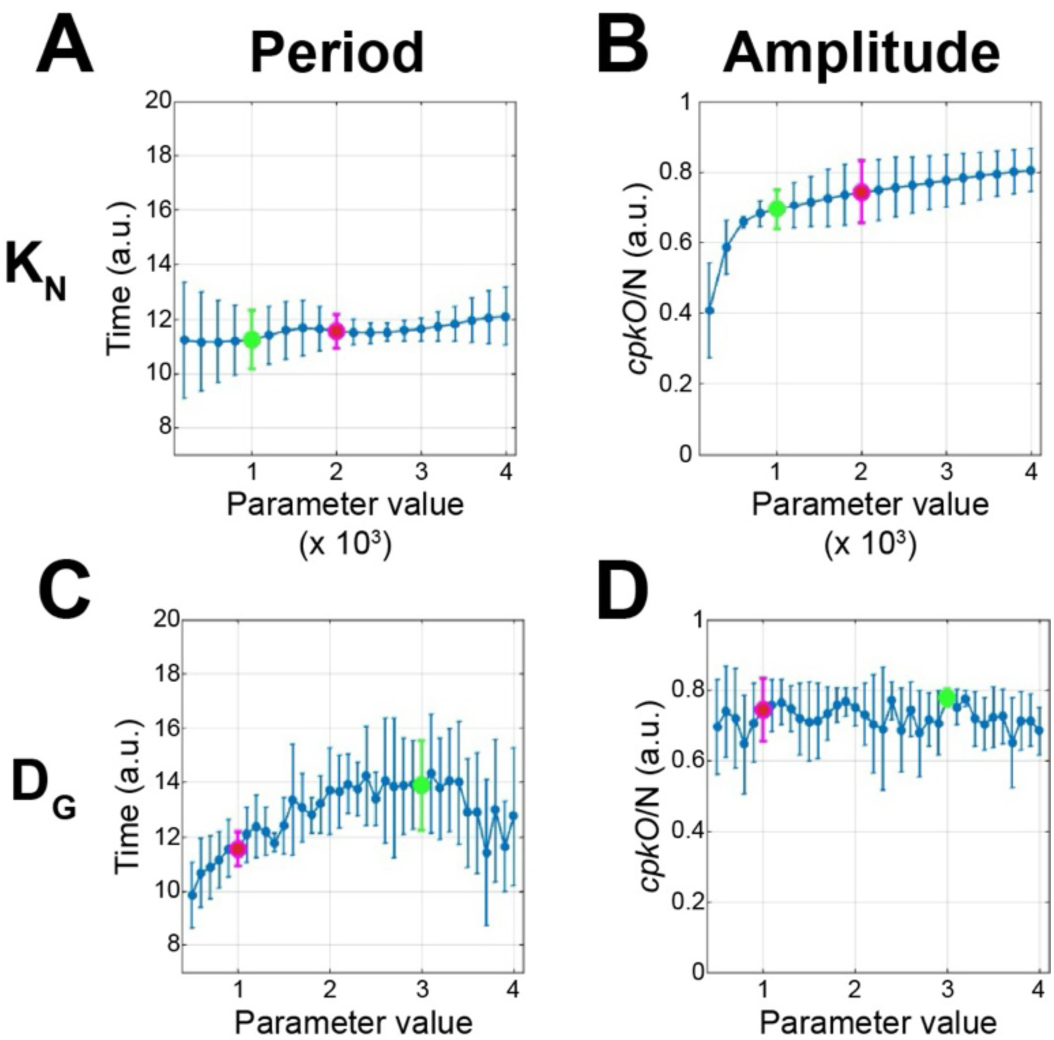
**Key model parameters may independently alter the periodicity or amplitude of simulated *cpkO* pulses**. Measured average period **(A, B)** or amplitude **(C, D**) of simulated *cpkO* expression oscillations at the growing colony edge when altering the diffusability of GBLs (D_G_) (A, C) or carrying capacity (K_N_) (B, D). Data points represent an average of oscillations that occurred during the duration of the simulation, with error bars representing SE. Data points highlighted in magenta indicate the nominal values of D_G_ or K_N_, Data points highlighted in green indicate the values that D_G_ or K_N_ were adjusted to for further analysis.

### Small molecule diffusabilty controls the periodicity of *cpkO* oscillations at the colony periphery

In our simulations, raising the diffusabilty (D_G_) to three times its normal value led to a period of 13.68 arbitrary time units compared to 11.62 at its normal value, an increase of 17.7%. This can be seen in normalized *cpkO* expression at the colony periphery which oscillated only three times rather than four within the same simulated time frame in this altered condition (Fig. 6A, 6B). Notably, changing the simulated carrying capacity (K_N_) did not alter the period (Fig. 6F, left).

**Fig. 6.**
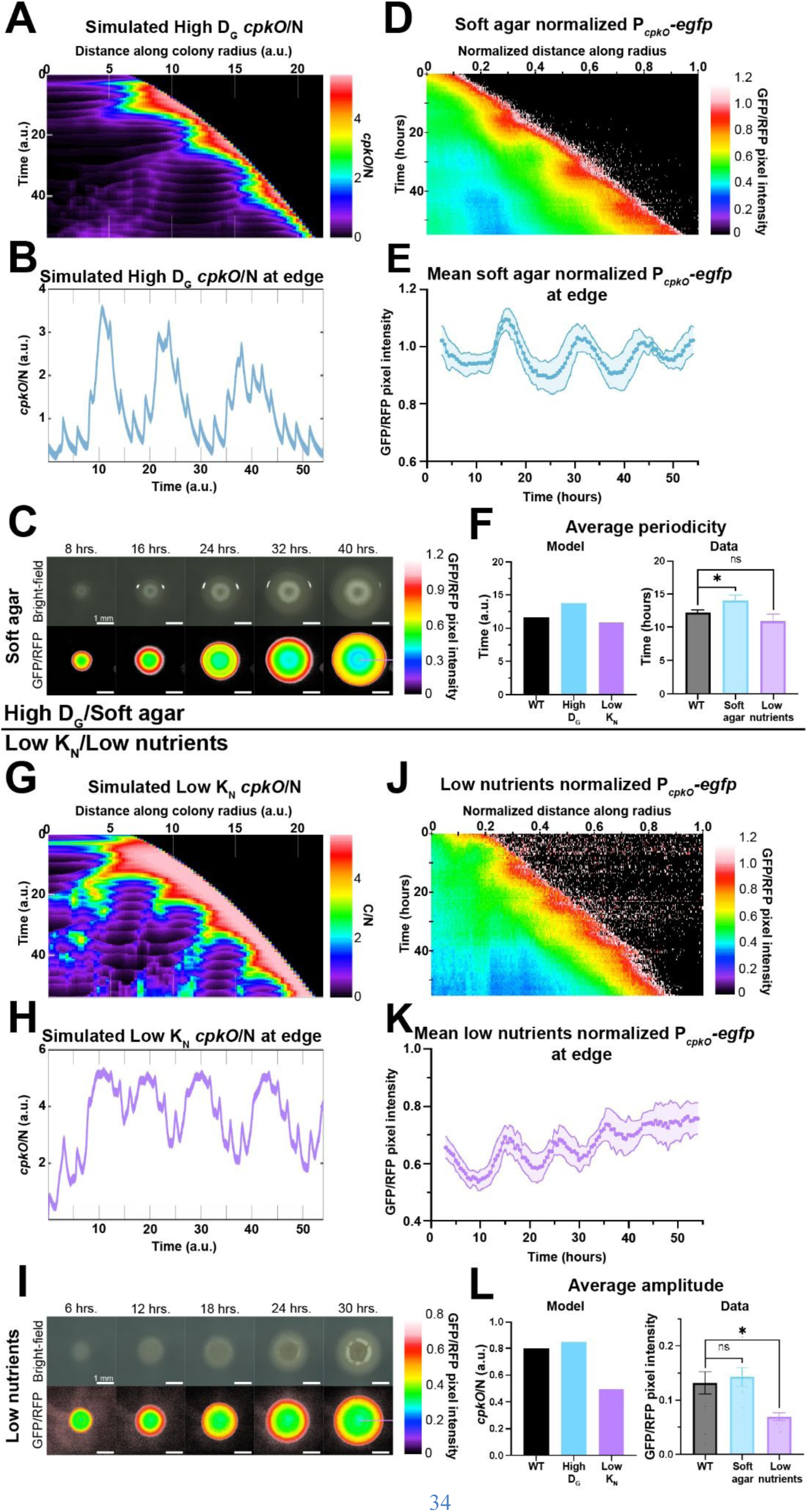
Environmental factors can modulate pulses of normalized *cpkO* expression. (A, **G)** Kymographs charting *cpkO*/N expression along the radius of a colony in a simulated environment with raised D_G_ (A) or lowered K_N_ (G). **(B, H)** Measured *cpkO*/N in a narrow band at the simulated colony edge for the raised D_G_ (B) or lowered K_N_ (H) condition. **(C, I)** Representative image time series of a real colony grown on soft agar (C) or nutrient-deprived (I) media. Images were curated from representative time-lapse movies. BF (top) and GFP/RFP (bottom) channels are shown. The color bar represents the LUT on which the GFP/RFP images are projected from. The mauve line indicates the axis on which the kymographs in D and I were built. Scale bars = 1 mm. **(D, J)** Kymographs charting P*_cpkO_-egfp*/P*_ermEp*_-mCherry* (GFP/RFP) expression along the radius of a colony grown on soft agar (D) or nutrient-deprived (J) media. **(E, K)** Fluorescence intensity plot that graphs GFP/RFP in a narrow band at the edge of colonies grown on soft agar (E) or nutrient-deprived media (K). The line denotes the mean of three measured colonies, with shading representing SE. SE for some data points may not be visible. **(F, L)** Measured average periodicity (F) or amplitude (L) of the WT and altered-environment oscillations observed in a simulated (left) and real (right) environment. Significant differences between means compared between indicated strains were determined using an unpaired two-tailed t-test (α = 0.05). “ns” indicates p > 0.05. “*” indicates p ≤ 0.05.

Guided by this model output, we next sought to perform a similar experiment with living *S. coelicolor* colonies. To do so, we grew colonies expressing the dual reporter system on ‘soft agar’ medium containing one-third the standard agar concentration (Fig. 6C). Lowering the agar concentration in the growth medium has been demonstrated to raise the diffusabilty of small molecules (45–47), and we confirmed this in our system by measuring point source diffusion of fluorescein on standard agar and soft agar (Fig. S7) (48, 49).

We found that colonies grown on soft agar exhibited ∼3.5 oscillations of normalized *cpkO*expression at their periphery within the 54 hr time frame (Fig. 6D, 6E) rather than the typical ∼4.5 oscillations observed on standard agar medium (Fig. 2C). Across three replicate colonies, growth on soft agar resulted in a significant increase in average periodicity to 14 hours compared to 12.21 hours on standard agar medium, an increase of 14.7% (Fig. 6F, right). These results indicate that the diffusabilty of GBLs directly impacts the periodicity of the *S. coelicolor* quorum oscillator.

### Carrying capacity controls the amplitude of *cpkO* oscillations at the colony periphery

In simulations, lowering the carrying capacity (K_N_ ) of the environment to one tenth of its normal value led to a wider peripheral band of normalized *cpkO* expression with less resolved oscillations (Fig. 6G). The amplitude of these pulses was diminished to 0.492 arbitrary units, as opposed to 0.799 arbitrary units in the standard model (Fig. 6H; 6L, left), a decrease of 38.4%.

To test this prediction, we grew *S. coelicolor* colonies on medium containing nutrients at a tenth of their standard concentration which effectively limits the carrying capacity. We then measured normalized *cpkO* expression, as done previously. Similar to the simulation, normalized *cpkO* expression oscillations were less resolved compared to those observed on standard medium (Fig. 6J, 6K). The amplitude of normalized *cpkO* oscillations across three colonies was lowered to 0.07 in this altered growth condition, a significant decrease of 46.2% from the amplitude of 0.13 observed under standard conditions (Fig. 6L, right). These results are aligned with the prediction that lowering the carrying capacity (and hence the number of cells in the colonies) results in a diminished oscillatory amplitude.

We note that in both simulations and experiments, changing either D_G_ or K_N_ altered either periodicity or amplitude respectively, but not both (Fig. 6F, 6L). We also note that altering the concentration of agar or nutrients did not significantly impact the rate of colony expansion over space, as is consistent with the model (Fig. S8). Taken together, these results demonstrate that environmental factors can predictably modulate different aspects of the behavior of *S. coelicolor* quorum oscillator.

## Discussion

The ability to generate periodic spatial patterns of gene expression is fundamental for organizing multicellular functions across diverse biological systems (50–56), yet few mechanisms capable of generating such spatial patterns have been characterized in bacteria (17). Here, we report a natural genetic oscillator in the multicellular bacterium *Streptomyces coelicolor* that integrates a quorum sensing gate into a repressilator-like topology. This quorum oscillator drives periodic spatial expression of a key regulator of antibiotic biosynthesis at the periphery of expanding *S. coelicolor* colonies (Fig. 2). Through a combination of mathematical modeling and genetic experiments, we show that this genetic circuit is sufficient to generate these periodic spatial patterns (Fig. 2), and mutants lacking circuit components fail to oscillate (Fig. 3, Fig. 4). Beyond this, the model predicted, and we experimentally verified, that the oscillator frequency and amplitude can be independently modulated by environmental factors, including diffusion limitation and carrying capacity, respectively (Fig. 5, Fig. 6). Thus, the *S. coelicolor* quorum oscillator enables collective, periodic spatial patterns to be structured in response to environmental conditions.

The *S. coelicolor* quorum oscillator does not directly parallel a classical repressilator. Rather, one step is replaced with a quorum-gated inhibitory connection (Fig. 1A, 1B). This topology ensures that progression through the circuit only occurs with adequate accumulation of the quorum signal. Inclusion of a quorum-sensing checkpoint into a repressilator-like circuit thus enables *S. coelicolor* colonies to synchronize oscillations as a collective across both space and time. This synchronization likely also limits the noise inherent in oscillations resulting from the original repressilator circuit (18–21).

The feasibility of using quorum control to produce collective, synchronized oscillations of gene expression has been demonstrated synthetically by using a simple genetic feedback loop (57). We note that in that system, it was necessary to trap the bacteria in a microfluidic context that enabled constant flow of the quorum signal away from the colony. In contrast, the natural circuit we describe here enables the emergence of reliably synchronized oscillations at the colony periphery, as cells grow into new territory where the quorum signal is absent. Thus, the *S. coelicolor* quorum oscillator provides a novel circuit topology for a variety of future synthetic systems where spatially organized periodic behavior is desired.

The observed oscillations of *cpkO* expression at the colony periphery may represent a unique strategy adopted by *S. coelicolor* with regard to antibiotic production (58–60). Why might *S. coelicolor* opt to make coelimycin antibiotics intermittently, rather than just turning antibiotic production on or off? Broadly, we speculate that synthesizing antibiotics at the colony periphery may be advantageous as cells expand into new space where competitors may be encountered. Oscillating production of coelimycins may save resources if pulsing antibiotic production is just as efficient as continuously producing it in a competitive context. Alternatively, this quorum oscillator may enable predictable, periodic reassessment of the local environment as new territory is explored, while potentially minimizing input from older, more remote cells where the local environment may be different. Further studies will be needed to assess the potential payoffs of these strategies.

Biological oscillators underlie periodic spatial pattern formation in diverse systems including vertebrate somitogenesis (9), arthropod body segmentation (10, 11), lateral root positioning in plants (12, 13), and sporulation zones in *Bacillus subtilis* biofilms (17). The *S. coelicolor* quorum oscillator shares a fundamental feature with these systems, namely, the coupling of oscillatory activity to an advancing growth front that converts temporal oscillations into periodic spatial patterns. However, the *S. coelicolor* quorum oscillator is markedly different from these systems in that its activity is inherently dependent on diffusible intercellular communication. The resulting architecture of this oscillator thus confers three emergent properties: synchronization of oscillatory activity across neighboring cells, dependence on collective cellular activity for progression through the regulatory cycle, and direct modulation in response to environmental conditions.

## Materials and Methods

### Strains and media

*S. coelicolor* M145 was obtained from the Traxler Lab at the University of California, Berkeley. This strain was designated as the wild-type (WT) strain for this study. For all experiments, *S. coelicolor* was grown on 20 mL of freshly poured R2YE agar at 30°C. For experiments with soft agar, the agar content of this media was lowered to a third of the normal concentration. For experiments with lowered nutrients, the glucose, casamino acids, and yeast extract were lowered to a tenth of their normal concentration. When performing conjugations, *S. coelicolor* was grown on mannitol soy (MS) agar supplemented with nalidixic acid (25 μg/mL) and either apramycin (20 μg/mL) or hygromycin (100 μg/mL) to select for successful exconjugants. *Escherichia coli* strains carrying plasmids conferring apramycin resistance were grown in LB Lennox supplemented with apramycin (20 μg/mL). Due to the effects of high salt concentration on hygromycin activity, *E. coli* strains carrying plasmids conferring hygromycin resistance were grown in LB Luria. The *E. coli* conjugation donor strain used in this study was ET12567/pUZ8002 (61, 62), which was grown in LB supplemented with kanamycin (50 μg/mL), chloramphenicol (25 μg/mL), and either apramycin or hygromycin at 37°C.

### Plasmid and strain construction

The Δ*scbA* and Δ*scbR2* mutants were constructed using CRISPomyces, a *Streptomyces*-optimized CRISPR/Cas-based gene editing system (63). This system was used to introduce single base-pair deletions to trigger early nonsense mutations 23 codons into the *scbA* CDS, and 48 codons into the *scbR2* CDS. The Δ*scbR* mutant was constructed using CRISPR-cBEST, a cytidine deaminase base editor which can replace a C:G base pair with an A:T base pair at a specific target site (64). This system was used to engineer an early nonsense mutation 51 codons into the *scbR* sequence. The online tool CRISPy-web was used to select appropriate protospacer targets for all three genes (65). These protospacers and their PAM sequences were aligned against the *S. coelicolor* genome using BLAST to minimize off-target binding. Curing the pCRISPomyces-2 and pCBEST vectors was accomplished by growing exconjugants without antibiotic selection to induce loss of the plasmid. Replica plating of cured strains on media with and without antibiotic selection was performed to confirm the successful loss of the plasmids.

The transcriptional reporter for *cpkO* was constructed in the pIJ8660 backbone and contains the *attP* integration site, φC31 *int* gene, EGFP, and apramycin resistance cassette (41, 66). To enhance the expression of EGFP in this plasmid and minimize any unexpected transcripts, the RNA element RiboJ was introduced into the pIJ8660 MCS using a Golden Gate Assembly (67). The promoter fragment for *cpkO* was amplified from M145 genomic DNA using Gibson Assembly-compatible primers and was cloned into pIJ8660 with a Gibson Assembly. The plasmid expressing mCherry is based on the pAV-1 backbone, and contains the *attP* integration site, VWB *int* gene, mCherry, RiboJ, and a hygromycin resistance cassette (66, 68). mCherry expression was placed under the control of *ermEp\**, a synthetic promoter designed for constitutive expression of downstream elements in *Streptomyces* (42, 66). The *ermEp\** sequence was obtained as a gBlock and cloned into pAV-1 with a Gibson Assembly. Promoterless EGFP and mCherry negative control vectors were obtained from the Traxler Lab (66).

### Microscopy

To get isolated colonies, a dilution series from a frozen *S. coelicolor* spore stock was plated. Isolated (< 10 colonies per plate) *S. coelicolor* colonies were incubated at 30°C for 42 hours before imaging. Time-lapse movies were recorded by capturing bright-field and fluorescence images every 30 minutes for 54 hours using a Zeiss Axio Zoom V16 microscope with a PlanNEOFLUAR Z 1.0x objective and an AxioCam 506 color camera. The light source used for fluorescence imaging was an Illuminator HXP 200C (metal halide), and the fluorescence filter sets used were FS 38HE (λ_excitation_ = 470 nm, λ_excitation_ = 525 nm) for EGFP and FS 43HE (λ_excitation_ = 550 nm, λ_excitation_ = 605 nm) for mCherry. The exposure time was set to 250 ms for the GFP channel, and to 1000 ms for the RFP channel. For the imaged P*_scbA_-egfp* and P*_bldN_-egfp* reporters, the GFP exposure time was increased to 1000 ms. To ensure an imaged colony remained at 30°C throughout the timecourse, we used a previously described protocol for constructing a temperature-controlled chamber under the microscope (66). Briefly, a temperature-controlled heated glass plate (Tokai Hit) was placed on top of a petri dish with the colony to be imaged. A temperature sensor (Tokai Hit) was sterilized and placed between the glass plate and the agar to monitor the temperature of this growth chamber. Representative individual GFP and RFP channels for all imaged strains and conditions are shown in Figure S9.

### Image processing

The time-lapse movies generated in this study were first processed following a previously described protocol (66). Briefly, mean autofluorescence from a negative control colony (*S. coelicolor* carrying promoter-less EGFP and mCherry constructs) was subtracted from a fluorescent colony for each frame of the movie in ImageJ. These movies were further processed in ImageJ to remove any remaining background fluorescence. The brightness and contrast of fluorescent time-lapse movies were further adjusted linearly in ImageJ for presentation purposes. To account for relatively high background in the GFP/RFP movies and images, the LUT was set to only display values from 0 to 1. This range was changed to 0-0.8 for the lower nutrient condition, and changed to 0-1.2 for the Δ*scbR2* and Δ*scbR* Δ*scbR2* strains, as well as the lower agar condition. These changes were made to adjust for lower and higher overall P*_cpkO_-egfp* intensity in these strains and conditions, respectively.

### Data visualization and analysis

All data were collected from time-lapse movies of the P*_cpkO_-egfp* reporter divided by a co-expressed P*_ermEp*_-mCherry* constitutive construct. This ratio ensures that any change in growth that is measured originates from genuine changes in reporter expression as opposed to changes in overall colony growth (69, 70). Kymographs were generated by first defining an ROI as the radius of an imaged colony at the final time point in the time course in ImageJ. EGFP reporter/mCherry constitutive expression was then measured along this radius for each frame of a time-lapse movie. Final kymographs were visualized in R. To account for high background that results from the EGFP/mCherry normalization, any value in an EGFP/mCherry kymograph above 1 (i.e. the EGFP pixel intensity is larger than the mCherry pixel intensity) was converted to 0. For colonies grown at a lower carrying capacity, this cutoff was lowered to 0.8 due to lower overall mCherry signal in this growth condition. Similarly, the cutoff was raised to 1.2 for Δ*scbR2* and Δ*scbR* Δ*scbR2* colonies, as well as those grown in the lower agar condition to compensate for overall more intense *cpkO* expression in this strain/growth condition. Representative EGFP/mCherry kymographs for each measured strain (n = 3) were chosen for main figures. All generated kymographs from separate and EGFP/mCherry normalized channels are shown in Figure S10-16.

Plots of EGFP/mCherry expression at the edge of the colony were generated by first defining an ROI as a band 40 pixels wide at the edge of an imaged growing colony for each frame of a time-lapse movie. Multiple band sizes were tested, and 40 pixels was selected as an optimal width (Figure S17). Mean fluorescence was then measured in this band at each time point. For the WT and low carrying capacity plots, each individual replicate was manually adjusted in time to align the phases of each oscillation across replicates. The data was then trimmed to ensure that each time point represented is in triplicate. Plots were visualized in GraphPad Prism.

The average amplitude for the observed pulses of normalized *cpkO* expression in both the model and data were calculated by measuring the expression at a trough then at the following peak to find the difference. Similarly, the average periodicity was calculated by measuring the time between each trough.

### Small molecule diffusion test

To assess how changing the agar concentration can change the diffusabilty of the QS signal, a simple experiment with fluorescein, a green fluorescent small molecule dye at a roughly similar molecular weight to a GBL, was performed (48, 71). 2.5 μL of fluorescein dye at a concentration of 2 μg/mL was spotted onto standard and soft agar R2YE plates, in triplicate (Figure S7A). After 30 minutes, these spots were imaged. Fluorescence intensity was measured across the diameter of the resulting fluorescent gradient discs to generate bell curves for both conditions (Figure S7B, C). The magnitude and distance across the disc to half-max fluorescence of these bell curves were calculated using Graphpad Prism. Statistical analyses were performed using two-tailed unpaired *t*-tests in Graphpad Prism to compare means between indicated conditions. The bell curve in the soft agar condition has a lower magnitude and higher distance to half-max, indicating that the shape of the bell curve has flattened, and that the fluorescein signal has diffused farther on soft agar compared to standard agar (Figure S7D, E).

### Mathematical model

The PDEs used to build the mathematical model are detailed in Figure S1. There are six PDEs incorporated into this model; five to spatiotemporally model each component of the *S. coelicolor* QS network, with an additional one to model spatiotemporal colony growth.

Using a standard logistic growth equation for the population coupled with a diffusion term, we capture the circularly/radially symmetric growth of the colony through the following equation: 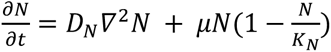, where *N* = *N*(*r*, *t*), *D_N_* represents the spatio-temporal diffusion coefficient of the cells in the environment, and *Δ*^2^*N* is the Laplacian operator. The second term in the equation above is the logistic growth model with growth rate *μ* and carrying capacity *K_N_*, which is assumed to be spatially uniform throughout the simulation. In this model, colony growth is decoupled from the QS network, and the population equation neglects cell death and other loss processes.

The following describes the spatio-temporal reaction-diffusion equations for each of the components of the genetic network depicted in Figure 1A. We note that these equations bear many similarities to others that simulate a standard repressilator circuit (Figure S18).

The variable A denotes *scbA*, G denotes GBLs, R denotes *scbR*, R2 denotes *scbR2*, finally, C denotes *cpkO*. Each of the genes in this circuit is characterized by three terms: (1) a standard positive, radially symmetric diffusion term, e.g. *D_A_Δ*^2^*A* for the dispersion of *scbA* in the environment; (2) a positive (potentially controlled) production rate, e.g. *α_G_A*, for the production of GBL; and (3) a negative diffusion/loss/degradation rate. e.g. −*δ_A_A* of *scbA*.

Having established that structure, we proceed with the description of the controlled production terms, which are dictated by the genetic circuit in terms of whether quantities are repressed or derepressed.

ScbA’s production occurs autonomously by each cell in the colony, however, its transcription is repressed by *scbR2*. Therefore, in its PDE, the production rate *α_A_* is multiplied by *N*, and by a repression Hill function of the form: 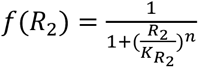, where *R* is the concentration of *scbR2*, *K_R_*_2*A*_ is the associated half-saturation constant, and *n* is the Hill coefficient that controls the strength of the repressor.

GBL production is promoted by ScbA and not repressed by any other genes. Therefore, in its associated PDE, the production rate *α_G_* is multiplied by *A*.

ScbR is autonomously produced by each cell unless it is self-repressed, which is quorum-controlled. Therefore, we add another term to the repression function to account for QS, denoted by *H*(*G*, *τ*), where *τ* is the QS threshold. Then, the production rate of *scbR*, *α_R_*, appears multiplied by *N* and by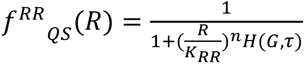

Similarly, *scbR2* is autonomously produced and repressed by ScbR, which only happens at quorum. Therefore, the production rate of *scbR2*, *α_R_*_2_, appears multiplied by *N* and by 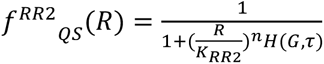

Finally, the production of *cpkO* is repressed by ScbR2 (not quorum-controlled) and ScbR (quorum-controlled). Therefore, the production rate of *cpkO*, *α_C_*, appears multiplied by *N* and by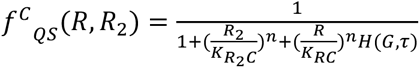

Lastly, we define the special QS activation function 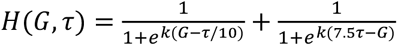, which in conjunction with the repression term operates as a band pass filter. That is, under quorum, *H*(*G*, *τ*) ≈ 0, the repression is turned off.

There are several parameters incorporated into these PDEs, all of which are shown in Table S1. These parameters were tuned using a hybrid approach based on reported relationships and experimental data generated here. For instance, the production of *cpkO* (α_C_) is substantially larger than the other production constants; this is supported by the native *cpkO* promoter generally being a very strong promoter (43). Additionally, the only diffusible component of this model is GBL’s; therefore, all diffusion constants except the one associated with GBLs (D_G_) were turned to zero. Lastly, the border of a *S. coelicolor* colony resembles a gently sloping mound as opposed to a sheer wall; therefore the eps parameter was set to a relatively low value to yield a realistic colony shape (44). After using reported relationships and biological truths to obtain the initial model parameters, we used the periodicity and amplitude of our generated experimental data to further refine key parameters (Δ_G_, α_G_, α_C_, α_R_, α_R2_, and Δ_C_). These key parameters were then further manually adjusted to yield clearly defined and resolved pulses of *cpkO*/N expression at the colony edge.

To explore the robustness of this model, we next performed a local sensitivity analysis in which a few key parameters listed above were substituted with many alternate values (Figure S19) (72). We then measured the periodicity and amplitude of the resulting *cpkO*/N pulses with these varied parameters. We demonstrate that oscillations with measurable amplitude and periodicity are still present at multiple values for several tested parameters (Fig. S19).

The signals produced by the mathematical model display lower frequency oscillations overlaid with high frequency ones. The high frequency component is likely a consequence of how the ring that defines the edge of the colony is propagating over time. The ring is also defined as a sharp region masking the portion of the space over which the signals are being averaged. That sharpness creates higher frequency components in the resulting average. To address this, we applied a low-pass filter on the signals, which was designed as follows. We used a third order butterworth filter with a cut off period Tc = 6h (i.e. the filter attenuates all spectral components corresponding to oscillations with periods less than or equal to 6h) (Fig. S20A). Equivalently, this eliminates all oscillations with frequency less than or equal to fc = 4.36 10^-5 Hz. This filtering approach smoothes the data to enable the automatic detection of peaks and valleys of simulated normalized *cpkO* pulses (Fig. S20B). Data that was smoothed by this filter was used for the sensitivity analysis.

Simulated kymographs and plots of *cpkO*/N at the colony edge were generated similarly to analogous ones generated for the data from real colonies. All figures pertaining to the model were generated and visualized in MATLAB. All code related to the mathematical model is publicly available on GitHub (https://github.com/TraxlerLab/Sco-quorum-oscillator-model).

## Competing interest statement

The authors declare no competing interest

## Acknowledgements

V.S. was funded in part by NIH grant 2T32GM132022-06 and the UC Berkeley Dissertation-Year Fellowship. M.M.V. was supported by startup funds from Florida State University. Work on genetic regulation of actinomycetes in the lab of M.F.T. is supported by NIH grant R35GM128849.

MV used Claude Fable 5 (Anthropic) to assist in writing and debugging the MATLAB code used for the numerical simulations and visualization of the quorum-sensing model.

All data are available in the main text, supplementary materials, and supplementary movies. All code is available in the online GitHub repository (https://github.com/TraxlerLab/Sco-quorum-oscillator-model)

## Notes

https://github.com/TraxlerLab/Sco-quorum-oscillator-model

### Competing Interest Statement

The authors have declared no competing interest.

## References

1. Li Z, Yang Q. 2018. Systems and synthetic biology approaches in understanding biological oscillators. Quant Biol 6:1–14.

2. Dunlap JC. 1999. Molecular bases for circadian clocks. Cell 96:271–290.

3. Satroutdinov AD, Kuriyama H, Kobayashi H. 1992. Oscillatory metabolism of Saccharomyces cerevisiae in continuous culture. FEMS Microbiol Lett 77:261–267.

4. Geva-Zatorsky N, Rosenfeld N, Itzkovitz S, Milo R, Sigal A, Dekel E, Yarnitzky T, Liron Y, Polak P, Lahav G, Alon U. 2006. Oscillations and variability in the p53 system. Mol Syst Biol 2:2006.0033.

5. Dupont G, Combettes L, Bird GS, Putney JW. 2011. Calcium oscillations. Cold Spring Harb Perspect Biol 3:a004226.

6. Kobayashi T, Mizuno H, Imayoshi I, Furusawa C, Shirahige K, Kageyama R. 2009. The cyclic gene Hes1 contributes to diverse differentiation responses of embryonic stem cells. Genes Dev 23:1870–1875.

7. Isomura A, Kageyama R. 2025. Progress in understanding the vertebrate segmentation clock. Nat Rev Genet 26:479–493.

8. Oates AC, Morelli LG, Ares S. 2012. Patterning embryos with oscillations: structure, function and dynamics of the vertebrate segmentation clock. Development 139:625–639.

9. Miao Y, Pourquié O. 2024. Cellular and molecular control of vertebrate somitogenesis. Nat Rev Mol Cell Biol 25:517–533.

10. Valentin G, Oates AC. 2013. Opening a can of centipedes: new insights into mechanisms of body segmentation. BMC Biol 11:116.

11. Diaz-Cuadros M, Pourquié O, El-Sherif E. 2021. Patterning with clocks and genetic cascades: Segmentation and regionalization of vertebrate versus insect body plans. PLoS Genet 17:e1009812.

12. Moreno-Risueno MA, Benfey PN. 2011. Time-based patterning in development: The role of oscillating gene expression. Transcription 2:124–129.

13. Perez-Garcia P, Serrano-Ron L, Moreno-Risueno MA. 2022. The nature of the root clock at single cell resolution: Principles of communication and similarities with plant and animal pulsatile and circadian mechanisms. Curr Opin Cell Biol 77:102102.

14. Ishiura M, Kutsuna S, Aoki S, Iwasaki H, Andersson CR, Tanabe A, Golden SS, Johnson CH, Kondo T. 1998. Expression of a gene cluster kaiABC as a circadian feedback process in cyanobacteria. Science 281:1519–1523.

15. Fang M, Partch CL, LiWang A, Golden SS. 2025. Prokaryotic Circadian Systems: Cyanobacteria and Beyond. Annu Rev Microbiol 79:523–545.

16. Lee CK, Schmidt WC, Webster SS, Chen JW, O’Toole GA, Wong GCL. 2022. Broadcasting of amplitude-and frequency-modulated c-di-GMP signals facilitates cooperative surface commitment in bacterial lineages. Proc Natl Acad Sci U S A 119:e2112226119.

17. Chou K-T, Lee D-YD, Chiou J-G, Galera-Laporta L, Ly S, Garcia-Ojalvo J, Süel GM. 2022. A segmentation clock patterns cellular differentiation in a bacterial biofilm. Cell 185:145–157.e13.

18. Elowitz MB, Leibler S. 2000. A synthetic oscillatory network of transcriptional regulators. Nature 403:335–338.

19. Potvin-Trottier L, Lord ND, Vinnicombe G, Paulsson J. 2016. Synchronous long-term oscillations in a synthetic gene circuit. Nature 538:514–517.

20. Zhang C, Tsoi R, You L. 2016. Addressing biological uncertainties in engineering gene circuits. Integr Biol (Camb) 8:456–464.

21. Purcell O, Savery NJ, Grierson CS, di Bernardo M. 2010. A comparative analysis of synthetic genetic oscillators. J R Soc Interface 7:1503–1524.

22. Waters CM, Bassler BL. 2005. Quorum sensing: cell-to-cell communication in bacteria. Annu Rev Cell Dev Biol 21:319–346.

23. Wang S, Garcia-Ojalvo J, Elowitz MB. 2022. Periodic spatial patterning with a single morphogen. Cell Syst 13:1033–1047.e7.

24. Hellen EH, Volkov E. 2017. Flexible dynamics of two quorum-sensing coupled repressilators. Phys Rev E 95:022408.

25. Garcia-Ojalvo J, Elowitz MB, Strogatz SH. 2004. Modeling a synthetic multicellular clock: repressilators coupled by quorum sensing. Proc Natl Acad Sci U S A 101:10955–10960.

26. Shum H, Balazs AC. 2017. Synthetic quorum sensing in model microcapsule colonies. Proc Natl Acad Sci U S A 114:8475–8480.

27. Fernández-Niño M, Giraldo D, Gomez-Porras JL, Dreyer I, González Barrios AF, Arevalo-Ferro C. 2017. A synthetic multi-cellular network of coupled self-sustained oscillators. PLoS One 12:e0180155.

28. Karig D, Martini KM, Lu T, DeLateur NA, Goldenfeld N, Weiss R. 2018. Stochastic Turing patterns in a synthetic bacterial population. Proc Natl Acad Sci U S A 115:6572–6577.

29. Tica J, Oliver Huidobro M, Zhu T, Wachter GKA, Pazuki RH, Bazzoli DG, Scholes NS, Tonello E, Siebert H, Stumpf MPH, Endres RG, Isalan M. 2024. A three-node Turing gene circuit forms periodic spatial patterns in bacteria. Cell Syst 15:1123–1132.e3.

30. Du P, Zhao H, Zhang H, Wang R, Huang J, Tian Y, Luo X, Luo X, Wang M, Xiang Y, Qian L, Chen Y, Tao Y, Lou C. 2020. De novo design of an intercellular signaling toolbox for multi-channel cell-cell communication and biological computation. Nat Commun 11:4226.

31. Bednarz B, Kotowska M, Pawlik KJ. 2019. Multi-level regulation of coelimycin synthesis in Streptomyces coelicolor A3(2). Appl Microbiol Biotechnol 103:6423–6434.

32. Takano E, Chakraburtty R, Nihira T, Yamada Y, Bibb MJ. 2001. A complex role for the gamma-butyrolactone SCB1 in regulating antibiotic production in Streptomyces coelicolor A3(2). Mol Microbiol 41:1015–1028.

33. Kato J, Funa N, Watanabe H, Ohnishi Y, Horinouchi S. 2007. Biosynthesis of gamma-butyrolactone autoregulators that switch on secondary metabolism and morphological development in Streptomyces. Proc Natl Acad Sci U S A 104:2378–2383.

34. Xu G, Wang J, Wang L, Tian X, Yang H, Fan K, Yang K, Tan H. 2010. “Pseudo” gamma-butyrolactone receptors respond to antibiotic signals to coordinate antibiotic biosynthesis. J Biol Chem 285:27440–27448.

35. Gottelt M, Kol S, Gomez-Escribano JP, Bibb M, Takano E. 2010. Deletion of a regulatory gene within the cpk gene cluster reveals novel antibacterial activity in Streptomyces coelicolor A3(2). Microbiology (Reading) 156:2343–2353.

36. Wang J, Wang W, Wang L, Zhang G, Fan K, Tan H, Yang K. 2011. A novel role of ’pseudo’γ-butyrolactone receptors in controlling γ-butyrolactone biosynthesis in Streptomyces. Mol Microbiol 82:236–250.

37. Li X, Wang J, Li S, Ji J, Wang W, Yang K. 2015. ScbR-and ScbR2-mediated signal transduction networks coordinate complex physiological responses in Streptomyces coelicolor. Sci Rep 5:14831.

38. Pawlik K, Kotowska M, Chater KF, Kuczek K, Takano E. 2007. A cryptic type I polyketide synthase (cpk) gene cluster in Streptomyces coelicolor A3(2). Arch Microbiol 187:87–99.

39. Bednarz B, Millan-Oropeza A, Kotowska M, Świat M, Quispe Haro JJ, Henry C, Pawlik K. 2021. Coelimycin Synthesis Activatory Proteins Are Key Regulators of Specialized Metabolism and Precursor Flux in Streptomyces coelicolor A3(2). Front Microbiol 12:616050.

40. Alon U. 2007. Network motifs: theory and experimental approaches. Nat Rev Genet 8:450– 461.

41. Sun J, Kelemen GH, Fernández-Abalos JM, Bibb MJ. 1999. Green fluorescent protein as a reporter for spatial and temporal gene expression in Streptomyces coelicolor A3(2). Microbiology (Reading) 145 (Pt 9):2221–2227.

42. Bibb MJ, Janssen GR, Ward JM. 1985. Cloning and analysis of the promoter region of the erythromycin resistance gene (ermE) of Streptomyces erythraeus. Gene 38:215–226.

43. Wang W, Li X, Wang J, Xiang S, Feng X, Yang K. 2013. An engineered strong promoter for streptomycetes. Appl Environ Microbiol 79:4484–4492.

44. Chater KF. 1998. Taking a genetic scalpel to the Streptomyces colony. Microbiology (Reading) 144:1465–1478.

45. Kümmerli R, Griffin AS, West SA, Buckling A, Harrison F. 2009. Viscous medium promotes cooperation in the pathogenic bacterium Pseudomonas aeruginosa. Proc Biol Sci 276:3531–3538.

46. Mund A, Diggle SP, Harrison F. 2017. The Fitness of Pseudomonas aeruginosa Quorum Sensing Signal Cheats Is Influenced by the Diffusivity of the Environment. mBio 8:e00353–17.

47. Ed-Daoui A, Snabre P. 2021. Poroviscoelasticity and compression-softening of agarose hydrogels. Rheol Acta 60:327–351.

48. Sjöback R, Nygren J, Kubista M. 1995. Absorption and fluorescence properties of fluorescein. Spectrochimica Acta Part A: Molecular and Biomolecular Spectroscopy 51:L7– L21.

49. Dilanji GE, Langebrake JB, De Leenheer P, Hagen SJ. 2012. Quorum activation at a distance: spatiotemporal patterns of gene regulation from diffusion of an autoinducer signal. J Am Chem Soc 134:5618–5626.

50. Aman AJ, Saunders LM, Carr AA, Srivatasan S, Eberhard C, Carrington B, Watkins-Chow D, Pavan WJ, Trapnell C, Parichy DM. 2023. Transcriptomic profiling of tissue environments critical for post-embryonic patterning and morphogenesis of zebrafish skin. Elife 12:RP86670.

51. Kondo S, Asal R. 1995. A reaction-diffusion wave on the skin of the marine angelfish Pomacanthus. Nature 376:765–768.

52. Sick S, Reinker S, Timmer J, Schlake T. 2006. WNT and DKK determine hair follicle spacing through a reaction-diffusion mechanism. Science 314:1447–1450.

53. Jung HS, Francis-West PH, Widelitz RB, Jiang TX, Ting-Berreth S, Tickle C, Wolpert L, Chuong CM. 1998. Local inhibitory action of BMPs and their relationships with activators in feather formation: implications for periodic patterning. Dev Biol 196:11–23.

54. Hu J, He L. 2008. Patterning mechanisms controlling digit development. J Genet Genomics 35:517–524.

55. Shoji H, Iwasa Y, Kondo S. 2003. Stripes, spots, or reversed spots in two-dimensional Turing systems. J Theor Biol 224:339–350.

56. Hentschel HGE, Glimm T, Glazier JA, Newman SA. 2004. Dynamical mechanisms for skeletal pattern formation in the vertebrate limb. Proc Biol Sci 271:1713–1722.

57. Danino T, Mondragón-Palomino O, Tsimring L, Hasty J. 2010. A synchronized quorum of genetic clocks. Nature 463:326–330.

58. Bystrykh LV, Fernández-Moreno MA, Herrema JK, Malpartida F, Hopwood DA, Dijkhuizen L. 1996. Production of actinorhodin-related “blue pigments” by Streptomyces coelicolor A3(2). J Bacteriol 178:2238–2244.

59. Tenconi E, Traxler MF, Hoebreck C, van Wezel GP, Rigali S. 2018. Production of Prodiginines Is Part of a Programmed Cell Death Process in Streptomyces coelicolor. Front Microbiol 9:1742.

60. Liu G, Chater KF, Chandra G, Niu G, Tan H. 2013. Molecular regulation of antibiotic biosynthesis in streptomyces. Microbiol Mol Biol Rev 77:112–143.

61. Paget MS, Chamberlin L, Atrih A, Foster SJ, Buttner MJ. 1999. Evidence that the extracytoplasmic function sigma factor sigmaE is required for normal cell wall structure in Streptomyces coelicolor A3(2). J Bacteriol 181:204–211.

62. Larcombe DE, Braes RE, Croxford JT, Wilson JW, Figurski DH, Hoskisson PA. 2024. Sequence and origin of the Streptomyces intergenetic-conjugation helper plasmid pUZ8002. Access Microbiol 6:000808.v3.

63. Cobb RE, Wang Y, Zhao H. 2015. High-efficiency multiplex genome editing of Streptomyces species using an engineered CRISPR/Cas system. ACS Synth Biol 4:723–728.

64. Tong Y, Whitford CM, Robertsen HL, Blin K, Jørgensen TS, Klitgaard AK, Gren T, Jiang X, Weber T, Lee SY. 2019. Highly efficient DSB-free base editing for streptomycetes with CRISPR-BEST. Proc Natl Acad Sci U S A 116:20366–20375.

65. Blin K, Pedersen LE, Weber T, Lee SY. 2016. CRISPy-web: An online resource to design sgRNAs for CRISPR applications. Synth Syst Biotechnol 1:118–121.

66. Zacharia VM, Ra Y, Sue C, Alcala E, Reaso JN, Ruzin SE, Traxler MF. 2021. Genetic Network Architecture and Environmental Cues Drive Spatial Organization of Phenotypic Division of Labor in Streptomyces coelicolor. mBio 12:e00794–21.

67. Clifton KP, Jones EM, Paudel S, Marken JP, Monette CE, Halleran AD, Epp L, Saha MS. 2018. The genetic insulator RiboJ increases expression of insulated genes. J Biol Eng 12:23.

68. Phelan RM, Sachs D, Petkiewicz SJ, Barajas JF, Blake-Hedges JM, Thompson MG, Reider Apel A, Rasor BJ, Katz L, Keasling JD. 2017. Development of Next Generation Synthetic Biology Tools for Use in Streptomyces venezuelae. ACS Synth Biol 6:159–166.

69. Repele A, Manu. 2019. Robust Normalization of Luciferase Reporter Data. Methods Protoc 2:62.

70. Su S, Bangar H, Saldanha R, Pemberton A, Aronow B, Dean GE, Lamkin TJ, Hassett DJ. 2014. Construction and characterization of stable, constitutively expressed, chromosomal green and red fluorescent transcriptional fusions in the select agents, Bacillus anthracis, Yersinia pestis, Burkholderia mallei, and Burkholderia pseudomallei. Microbiologyopen 3:610–629.

71. Takano E, Nihira T, Hara Y, Jones JJ, Gershater CJ, Yamada Y, Bibb M. 2000. Purification and structural determination of SCB1, a gamma-butyrolactone that elicits antibiotic production in Streptomyces coelicolor A3(2). J Biol Chem 275:11010–11016.

72. Hamby DM. 1994. A review of techniques for parameter sensitivity analysis of environmental models. Environ Monit Assess 32:135–154.

73. Bentley SD, Chater KF, Cerdeño-Tárraga A-M, Challis GL, Thomson NR, James KD, Harris DE, Quail MA, Kieser H, Harper D, Bateman A, Brown S, Chandra G, Chen CW, Collins M, Cronin A, Fraser A, Goble A, Hidalgo J, Hornsby T, Howarth S, Huang C-H, Kieser T, Larke L, Murphy L, Oliver K, O’Neil S, Rabbinowitsch E, Rajandream M-A, Rutherford K, Rutter S, Seeger K, Saunders D, Sharp S, Squares R, Squares S, Taylor K, Warren T, Wietzorrek A, Woodward J, Barrell BG, Parkhill J, Hopwood DA. 2002. Complete genome sequence of the model actinomycete Streptomyces coelicolor A3(2). Nature 417:141–147.

