## Supplementary Figures for "Periodic spatial patterning of gene expression via a natural quorum-synchronized oscillator"

**<u>Contents:</u>**

Supplementary Figure 1: **Partial differential equations (PDEs) used to construct the mathematical model**

Supplemental Figure 2: **Expression of a gene not directly QS-controlled (*bldN*) does not pulse at the colony edge**

Supplemental Figure 3: ***scbA* has relatively low expression in WT *S. coelicolor* colonies**

Supplemental Figure 4: **Normalized *cpkO* expression at the edge of virtual and real *S. coelicolor* colonies**

Supplemental Figure 5: **Kymographs of simulated normalized *cpkO* expression in *ΔscbA* colonies with varied repressive strengths of ScbR on itself and *scbR2*, and *cpkO***

Supplemental Figure 6: **Kymographs of simulated *cpkO* expression in *ΔscbR* colonies with varied repressive strengths of ScbR2 on *cpkO***

Supplemental Figure 7: **Small molecules diffuse farther on soft agar compared to standard agar**

Supplemental Figure 8: **Quantifying radial expansion of colonies grown on standard and altered media**

Supplemental Figure 9: **Representative GFP and RFP time series for all imaged strains and conditions**

Supplemental Figure 10: **GFP, RFP, and GFP/RFP kymographs for replicates of WT colonies**

Supplemental Figure 11: **GFP, RFP, and GFP/RFP kymographs for replicates of Δ*scbA* colonies**

Supplemental Figure 12: **GFP, RFP, and GFP/RFP kymographs for replicates of Δ*scbR* colonies**.

Supplemental Figure 13: **GFP, RFP, and GFP/RFP kymographs for replicates of Δ*scbR2* colonies**

Supplemental Figure 14: **GFP, RFP, and GFP/RFP kymographs for replicates of Δ*scbR* Δ*scbR2* colonies.**

Supplemental Figure 15: **GFP, RFP, and GFP/RFP kymographs for replicates of WT colonies grown on soft agar**

Supplemental Figure 16: **GFP, RFP, and GFP/RFP kymographs for replicates of WT colonies grown on nutrient-deprived media**

Supplemental Figure 17: **ROI optimization for plots of normalized *cpkO* expression at the edge of a colony**

Supplemental Figure 18: **A mathematical model of a standard repressilator circuit.**

Supplemental Figure 19: **Sensitivity analysis on several model parameters**

Supplemental Figure 20: **Filtering *in silico* data to resolve high-frequency oscillations**

Table 1: **List of parameters incorporated into the model**

Table 2: **List of strains used in this study**

Movie 1: **Fluorescence time-lapse movie of $P_{cpkO}$-*egfp*/$P_{ermEp*}$-*mCherry* (GFP/RFP) expression in a representative WT colony.**

Movie 2: **Simulated *cpkO*/N in a virtual colony**

$$\frac{\partial N}{\partial t} = D_N \nabla^2 N + \mu N \left(1 - \frac{N}{K_N}\right)$$

$$\frac{\partial A}{\partial t} = D_A \nabla^2 A + \alpha_A \cdot \frac{N}{1 + \left(\frac{R_2}{K_{R_2 A}}\right)^n} - \delta_A A$$

$$\frac{\partial G}{\partial t} = D_G \nabla^2 G + \alpha_G A - \delta_G G$$

$$\frac{\partial R}{\partial t} = D_R \nabla^2 R + \alpha_R \cdot \frac{N}{1 + \left(\frac{R}{K_{RR}}\right)^n \cdot H(G,\tau)} - \delta_R R$$

$$\frac{\partial R_2}{\partial t} = D_{R_2} \nabla^2 R_2 + \alpha_{R_2} \cdot \frac{N}{1 + \left(\frac{R}{K_{RR2}}\right)^n \cdot H(G,\tau)} - \delta_{R_2} R_2$$

$$\frac{\partial C}{\partial t} = D_C \nabla^2 C + \alpha_C \cdot \frac{N}{1 + \left(\frac{R_2}{K_{R_2 C}}\right)^n + \left(\frac{R}{K_{RC}}\right)^n \cdot H(G,\tau)} - \delta_C C$$

**Figure S1:** Partial differential equations (PDEs) used to construct the mathematical model. This model consists of five differential equations that simulate the expression of each component of the *S. coelicolor* circuit over space and time, and one equation to simulate spatiotemporal growth of the virtual colony. Full details regarding the model can be found in the Materials & Methods, and a full list of parameters incorporated into the model can be found in Supplementary Table 1.

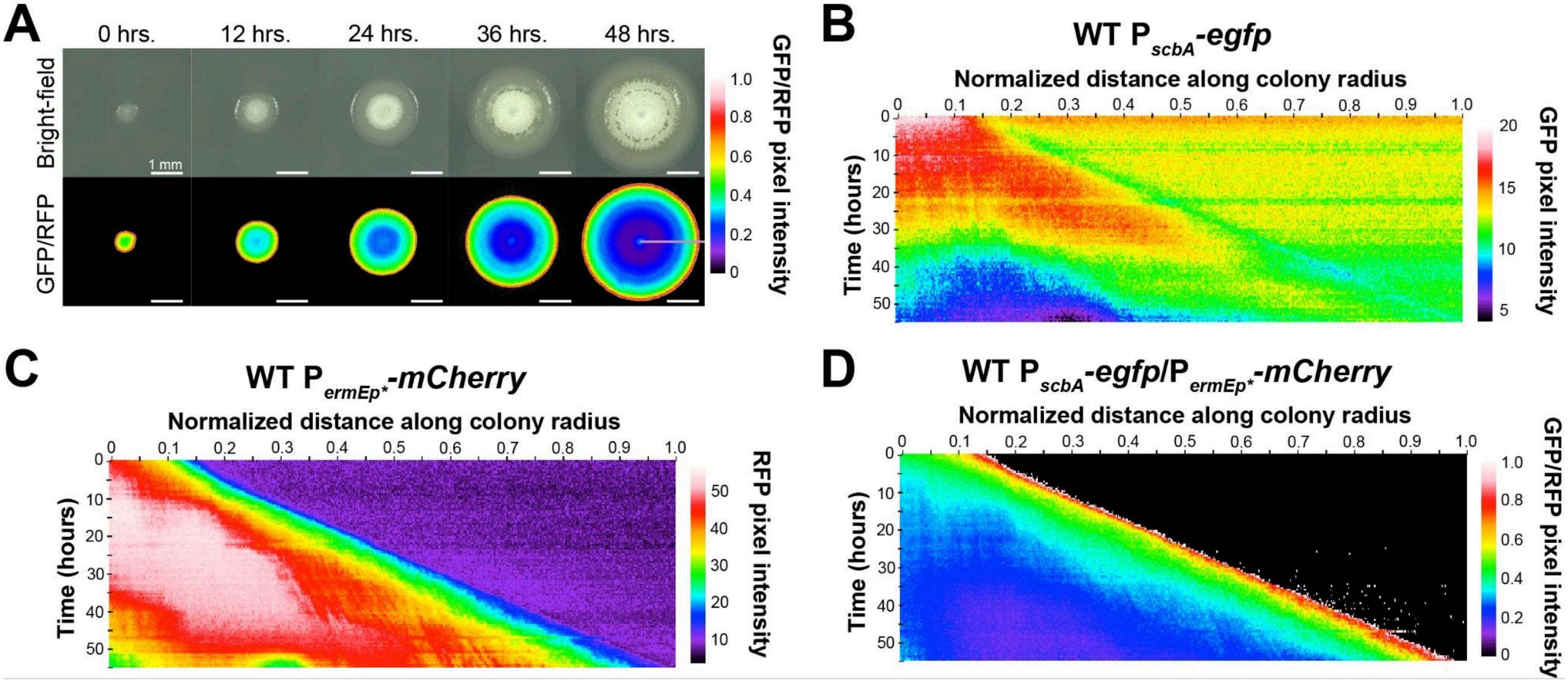


**Figure S2: Expression of a gene not directly QS-controlled (*bldN*) does not pulse at the colony edge. (A)** Representative BF and GFP/RFP images showing growth of a WT colony (top) and $P_{bldN}$-*egfp*/$P_{ermEp*}$-*mCherry* expression in that colony (bottom) over 48 hours. The mauve line in the last GFP/RFP image shows the radius on which the kymographs in B, C, and D are built. Scale bar - 1 mm. **(B,C,D)** Kymographs charting $P_{bldN}$-*egfp* (B), $P_{ermEp*}$-*mCherry* (C), and $P_{bldN}$-*egfp*/$P_{ermEp*}$-*mCherry* expression (D).

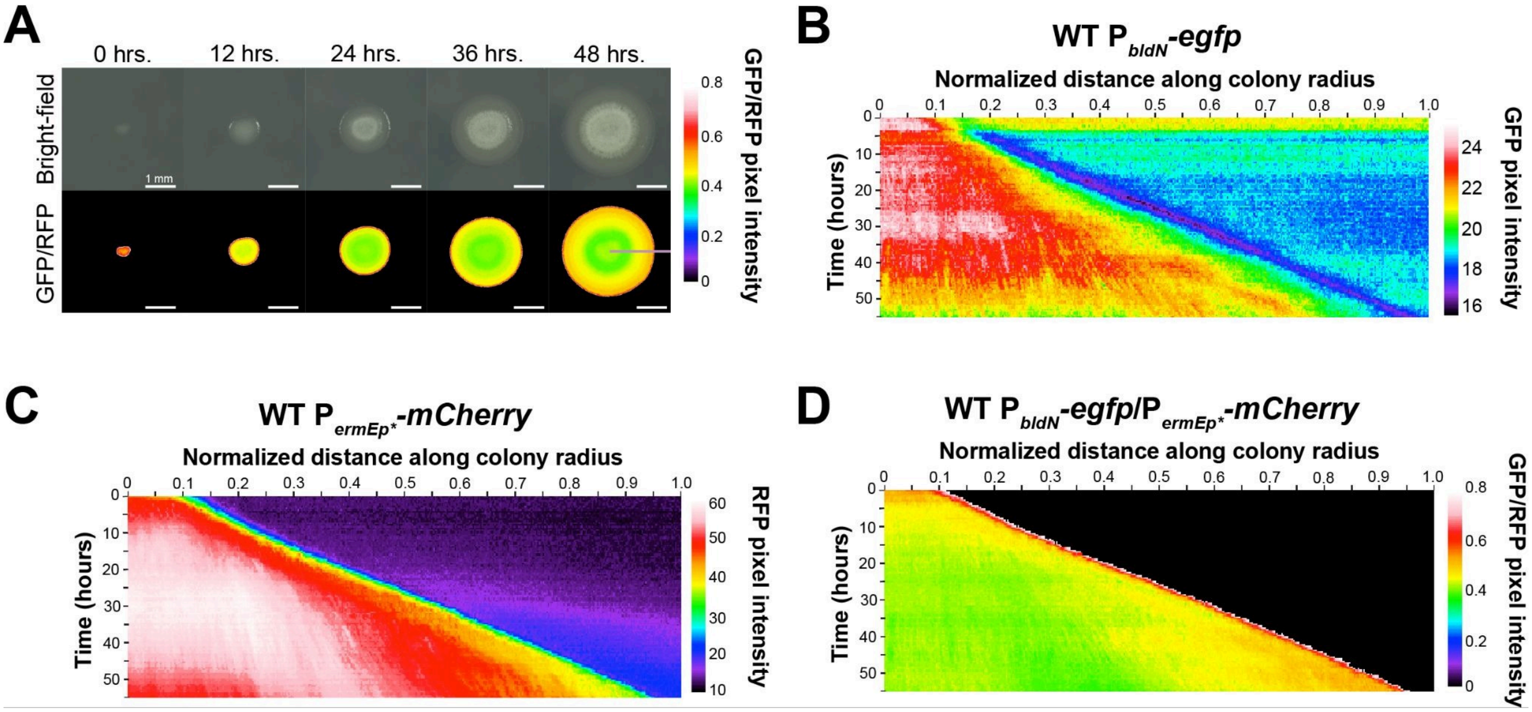


**Figure S3: *scbA* has relatively low expression in WT *S. coelicolor* colonies. (A)** Representative BF and GFP/RFP images showing growth of a WT colony (top) and $P_{scbA}$-*egfp*/$P_{ermEp*}$-*mCherry* expression in that colony (bottom) over 48 hours. The mauve line in the last GFP/RFP image shows the radius on which the kymographs in B, C, and D are built. Scale bar - 1 mm. **(B,C,D)** Kymographs charting $P_{scbA}$-*egfp* (B), $P_{ermEp*}$-*mCherry* (C), and $P_{scbA}$-*egfp*/$P_{ermEp*}$-*mCherry* expression (D). Normalized *scbA* expression appears as only a very thin line at the colony periphery.

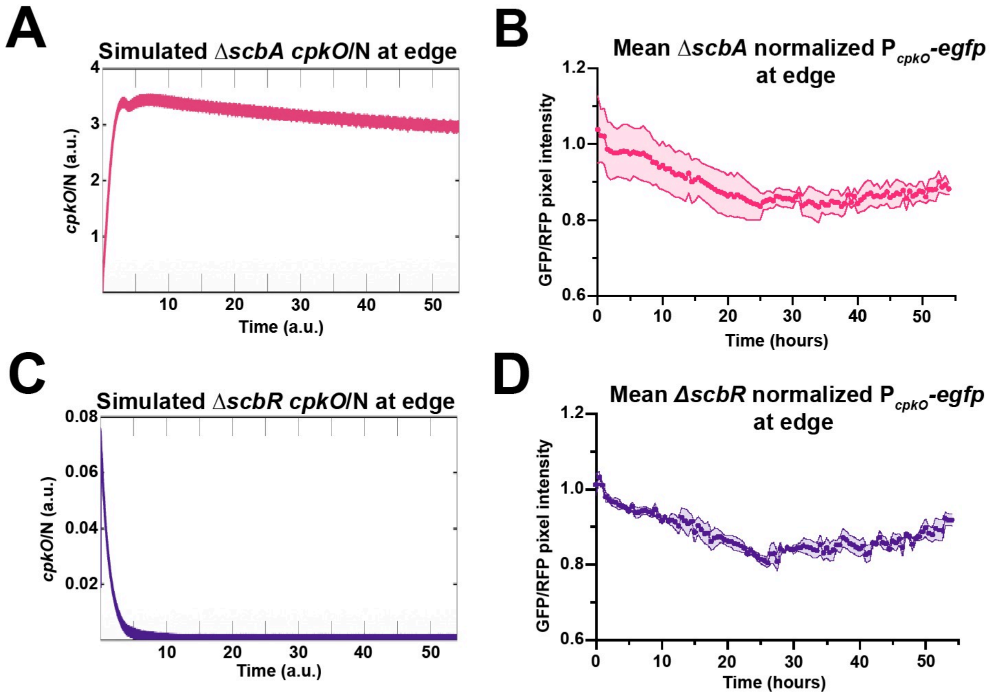


**Figure S4: Normalized *cpkO* expression at the edge of virtual and real *S. coelicolor* colonies. (A, C)** Measured *cpkO*/N in a narrow band at the simulated Δ*scbA* (A) or Δ*scbR* (C) colony edge. **(B, D)** Fluorescence intensity plot that graphs GFP/RFP in a narrow band at the edge of Δ*scbA* (B) or Δ*scbR* (D) colonies. The line denotes the mean of three measured colonies, with shading representing SE. SE for some data points may not be visible.

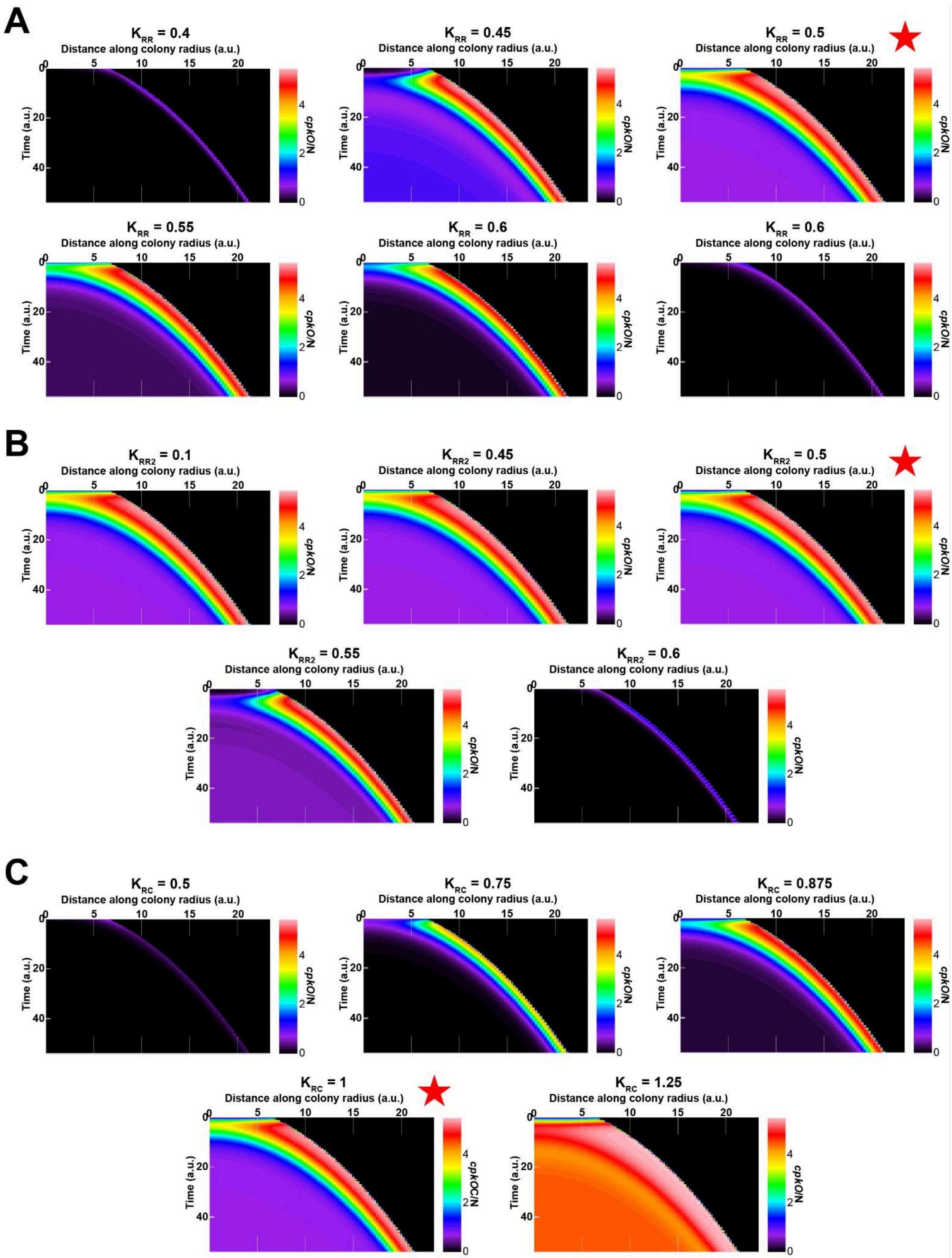
A
K_RR = 0.4
K_RR = 0.45
K_RR = 0.5
K_RR = 0.55
K_RR = 0.6
K_RR = 0.6
B
K_RR2 = 0.1
K_RR2 = 0.45
K_RR2 = 0.5
K_RR2 = 0.55
K_RR2 = 0.6
C
K_RC = 0.5
K_RC = 0.75
K_RC = 0.875
K_RC = 1
K_RC = 1.25
Distance along colony radius (a.u.)
Time (a.u.)
cpkO/N

**Figure S5: Kymographs of simulated normalized *cpkO* expression in ∆*scbA* colonies with varied repressive strengths of ScbR on itself and *scbR2*, and *cpkO*.** Multiple tested parameters values for $K_{RR}$ **(A)**, $K_{RR2}$ **(B)**, or $K_{RC}$ **(C)** and how they influence normalized *cpkO* expression in a simulated ∆*scbA* colony. All kymographs chart normalized *cpkO* expression along the radius of a simulated colony. Red stars indicate the standard value of the parameter.

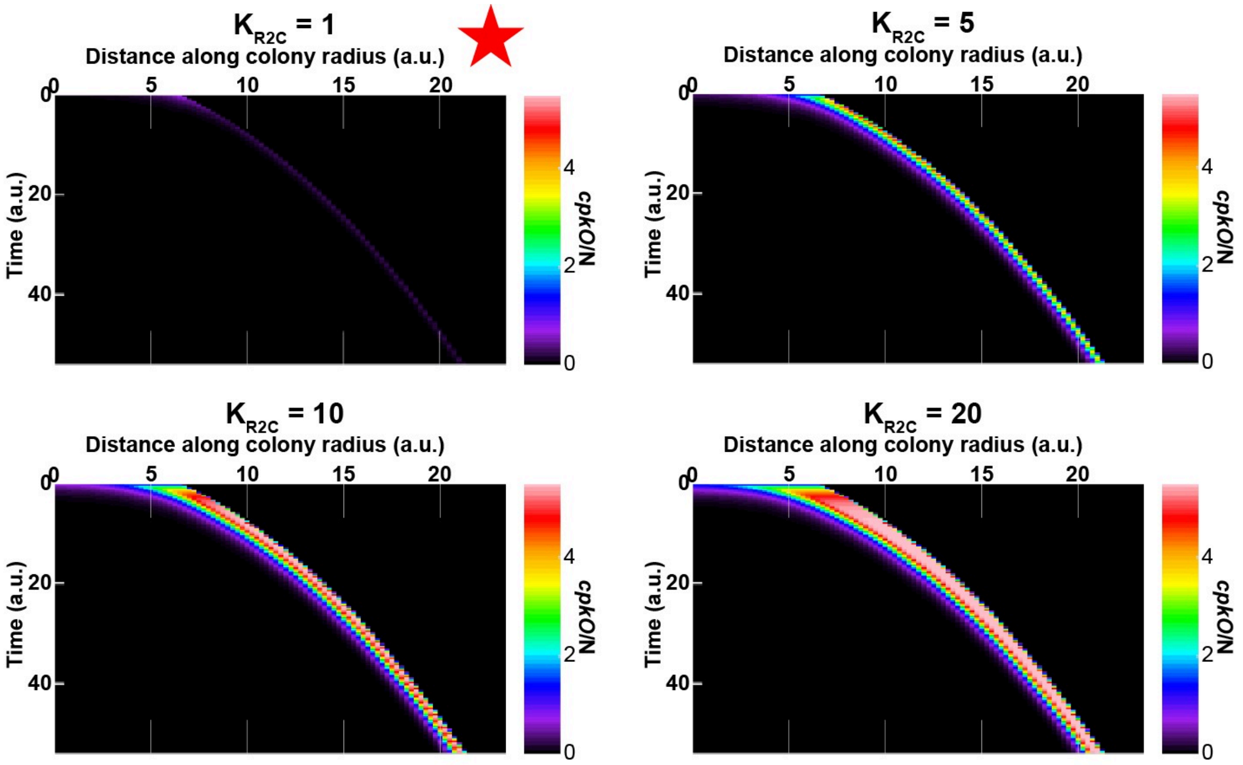


**Figure S6: Kymographs of simulated *cpkO* expression in ∆*scbR* colonies with varied repressive strengths of ScbR2 on *cpkO*.** Multiple tested parameters values for $K_{R2C}$ and how they influence normalized *cpkO* expression in a simulated ∆*scbR* colony. All kymographs chart normalized *cpkO* expression along the radius of a simulated colony. The red star indicates the standard value of the parameter.

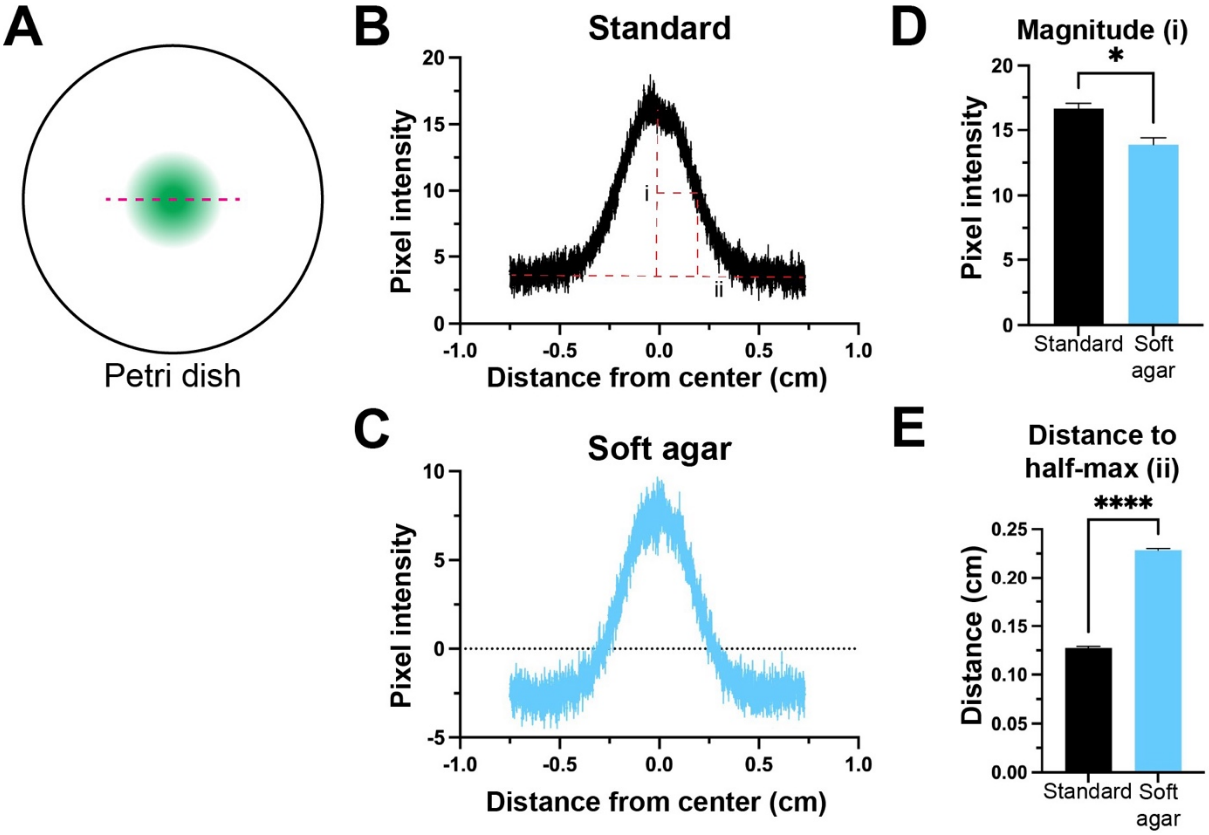


**Figure S7: Small molecules diffuse farther on soft agar compared to standard agar. (A)** mockup of how fluorescein would diffuse in agar from a drop on a petri dish. The gray dashed line indicates the axis on which fluorescence intensity was measured. **(B, C)** Fluorescence intensity after background subtraction along the measured axis in (A) for a drop of fluorescein dye on standard (B) or soft (C) agar. The line represents the mean of three data points, with error bars representing SE. (B) Red dashed lines indicate how the magnitude (i) and distance to half-max fluorescence (ii) were calculated. (D) Measured magnitude **(D)** and distance to half-max **(E)** of the curves in (B) and (C). Significant differences between means compared between indicated strains were determined using an unpaired two-tailed t-test (α = 0.05). "*" indicates $p \leq 0.05$ and "****" indicates $p \leq 0.0001$.

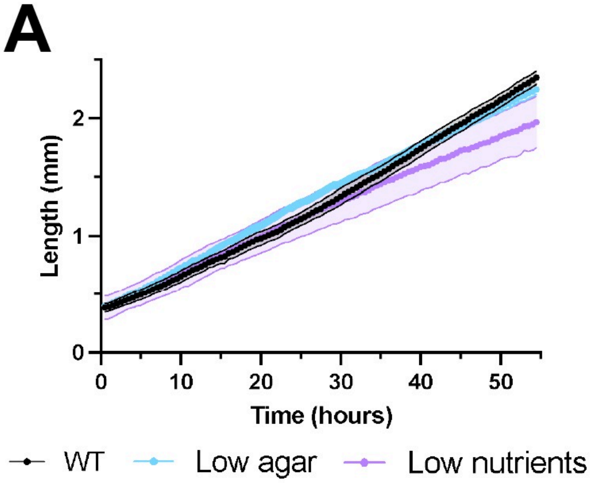


**Figure S8: Quantifying radial expansion of colonies grown on standard and altered media. (A)** The radius of colonies grown on standard media, soft agar media, or nutrient-deprived media over time. Points represent the mean of three measured colonies, and shading represents SE. **(B)** Rate of radius expansion of the colonies measured in (A). Bars represent the mean of three colonies, and error bars represent SE. Significant differences between means compared between indicated strains were determined using an unpaired two-tailed t-test ($\alpha$ = 0.05). “ns” indicates $p > 0.05$.

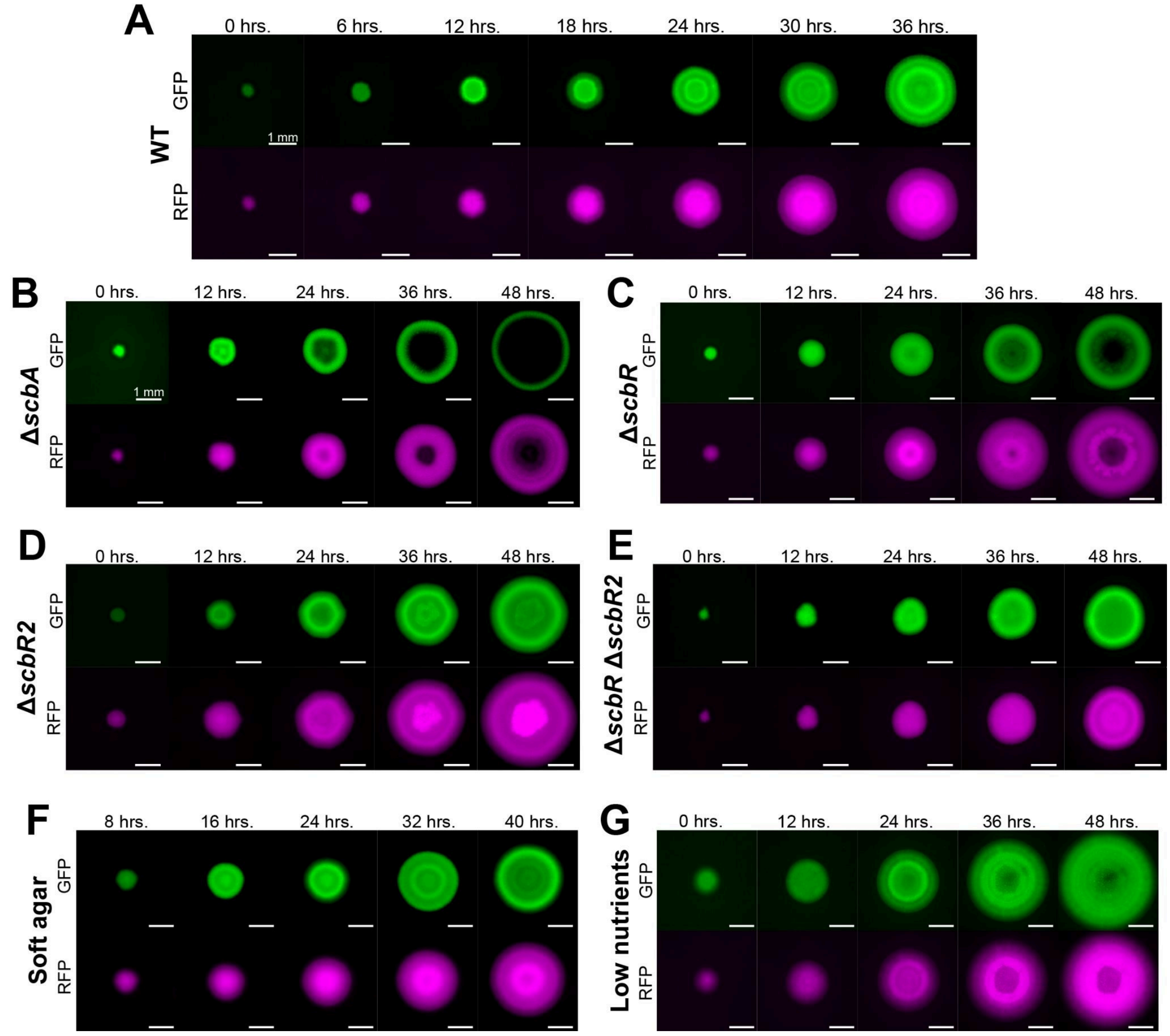


**Figure S9: Representative GFP and RFP time series for all imaged strains and conditions.** Non-normalized GFP and RFP channels for representative replicates for all strains/conditions observed in this study. Images were scaled linearly for presentation purposes. Scale bars = 1 mm.

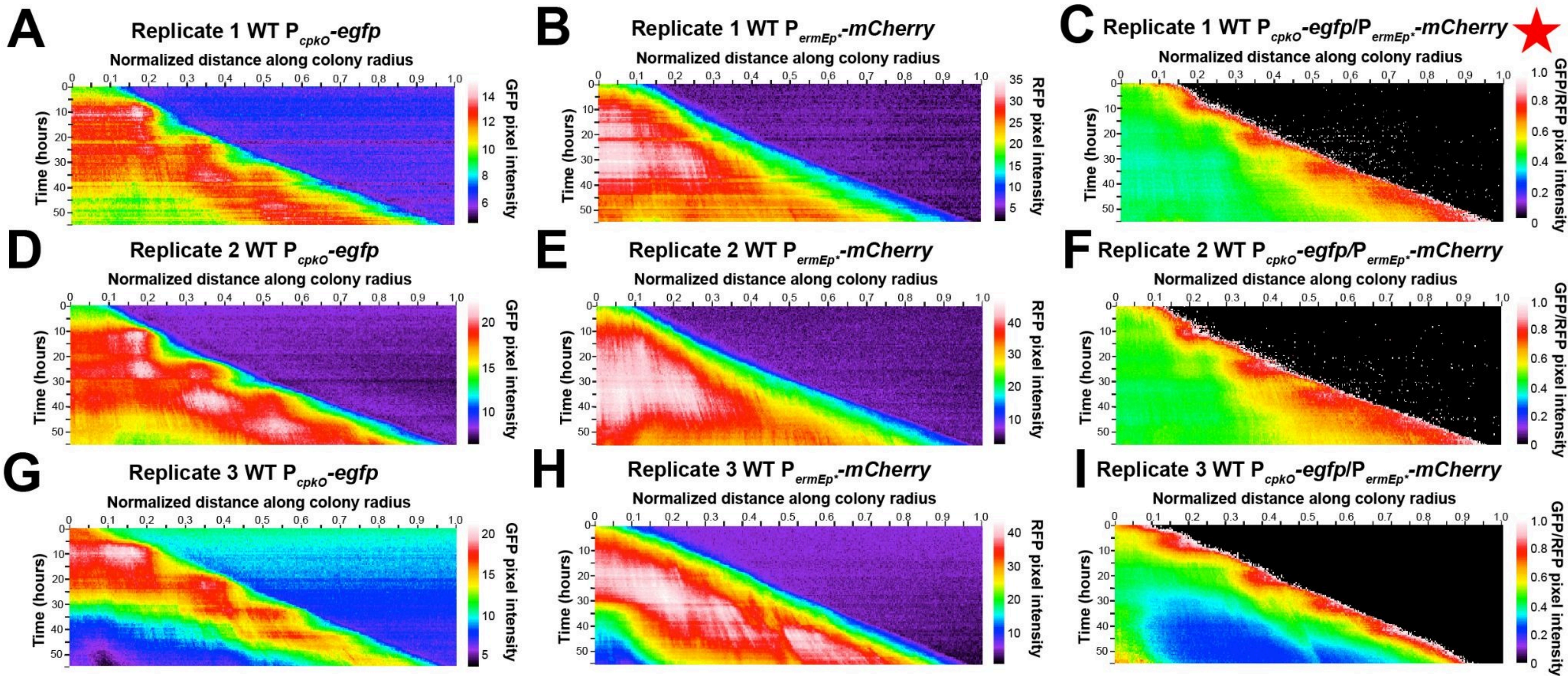


**Figure S10: GFP, RFP, and GFP/RFP kymographs for replicates of WT colonies**. Individual GFP **(A, D, G)**, RFP **(B, E, H)** , and GFP/RFP **(C, F, I)** kymographs for three measured WT *S. coelicolor* colonies (A-C, D-F, G-I). The star indicates the representative kymograph shown in Figure 2B.

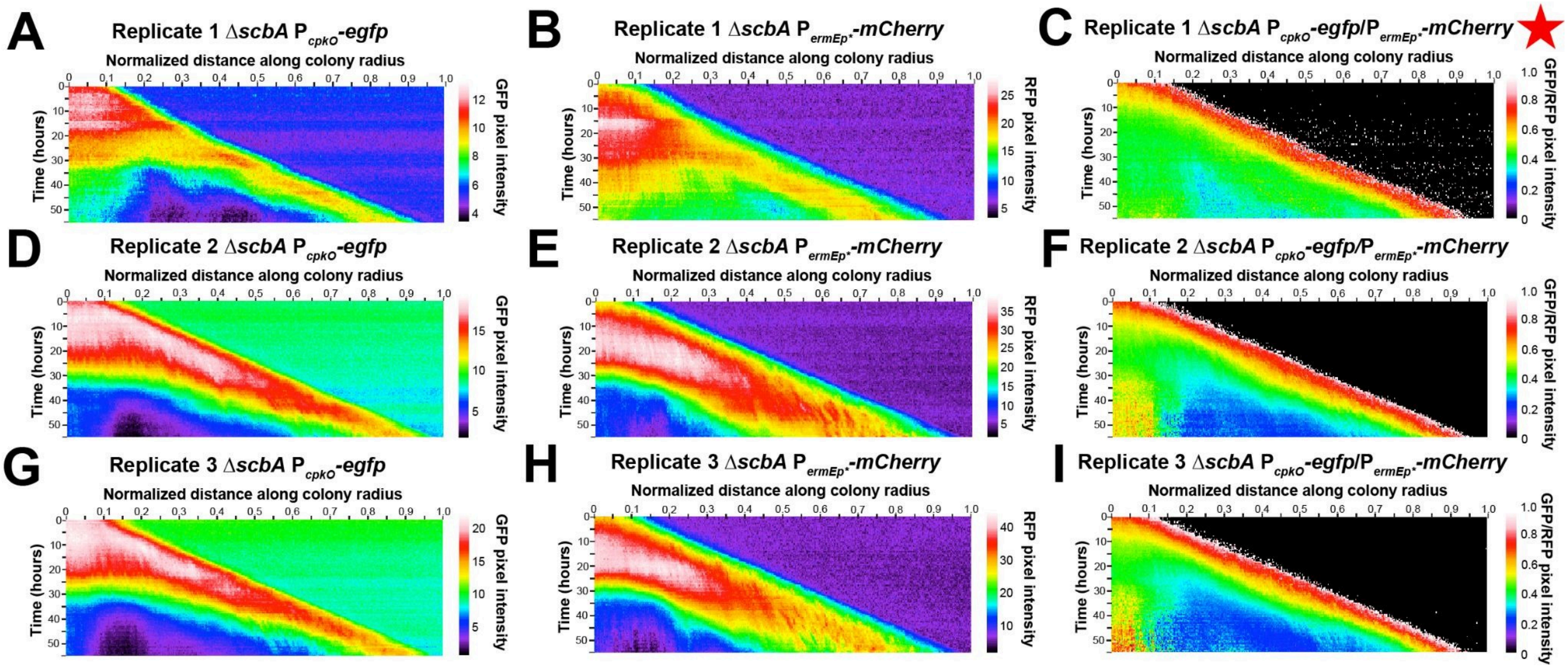


**Figure S11: GFP, RFP, and GFP/RFP kymographs for replicates of Δ*scbA* colonies**. Individual GFP **(A, D, G)**, RFP **(B, E, H)** , and GFP/RFP **(C, F, I)** kymographs for three measured Δ*scbA* colonies (A-C, D-F, G-I). The star indicates the representative kymograph shown in Figure 3D.

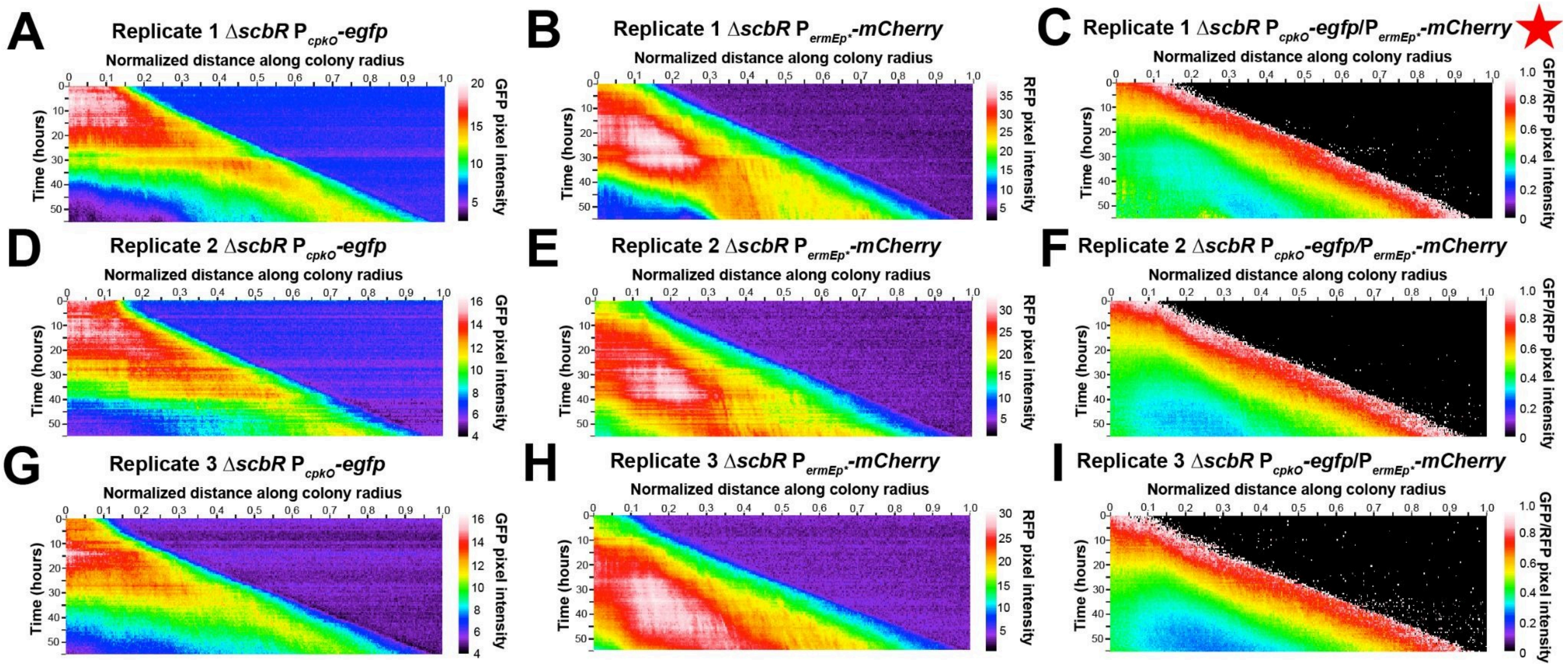


**Figure S12: GFP, RFP, and GFP/RFP kymographs for replicates of Δ*scbR* colonies**. Individual GFP **(A, D, G)**, RFP **(B, E, H)**, and GFP/RFP **(C, F, I)** kymographs for three measured Δ*scbR* colonies (A-C, D-F, G-I). The star indicates the representative kymograph shown in Figure 3I.

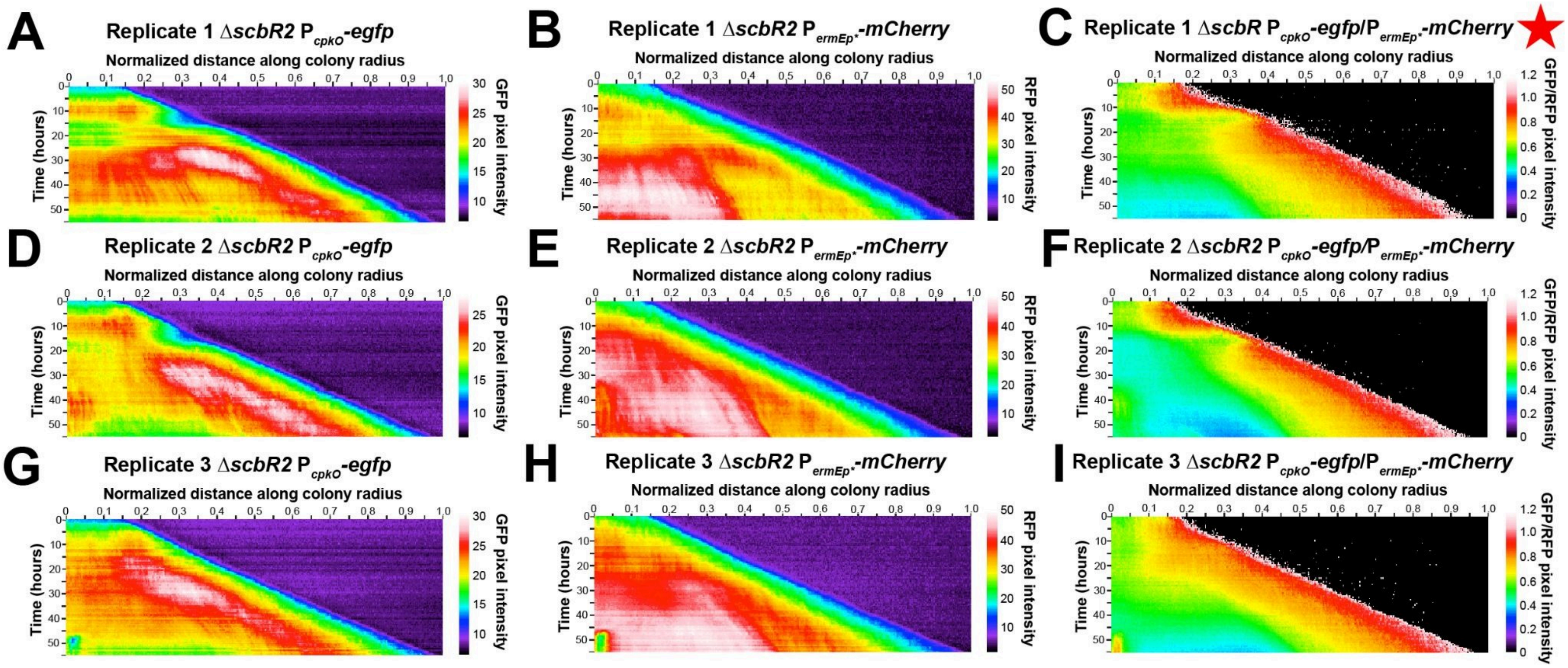


**Figure S13: GFP, RFP, and GFP/RFP kymographs for replicates of Δ*scbR2* colonies**. Individual GFP **(A, D, G)**, RFP **(B, E, H)** , and GFP/RFP **(C, F, I)** kymographs for three measured Δ*scbR2* colonies (A-C, D-F, G-I). The star indicates the representative kymograph shown in Figure 4D.

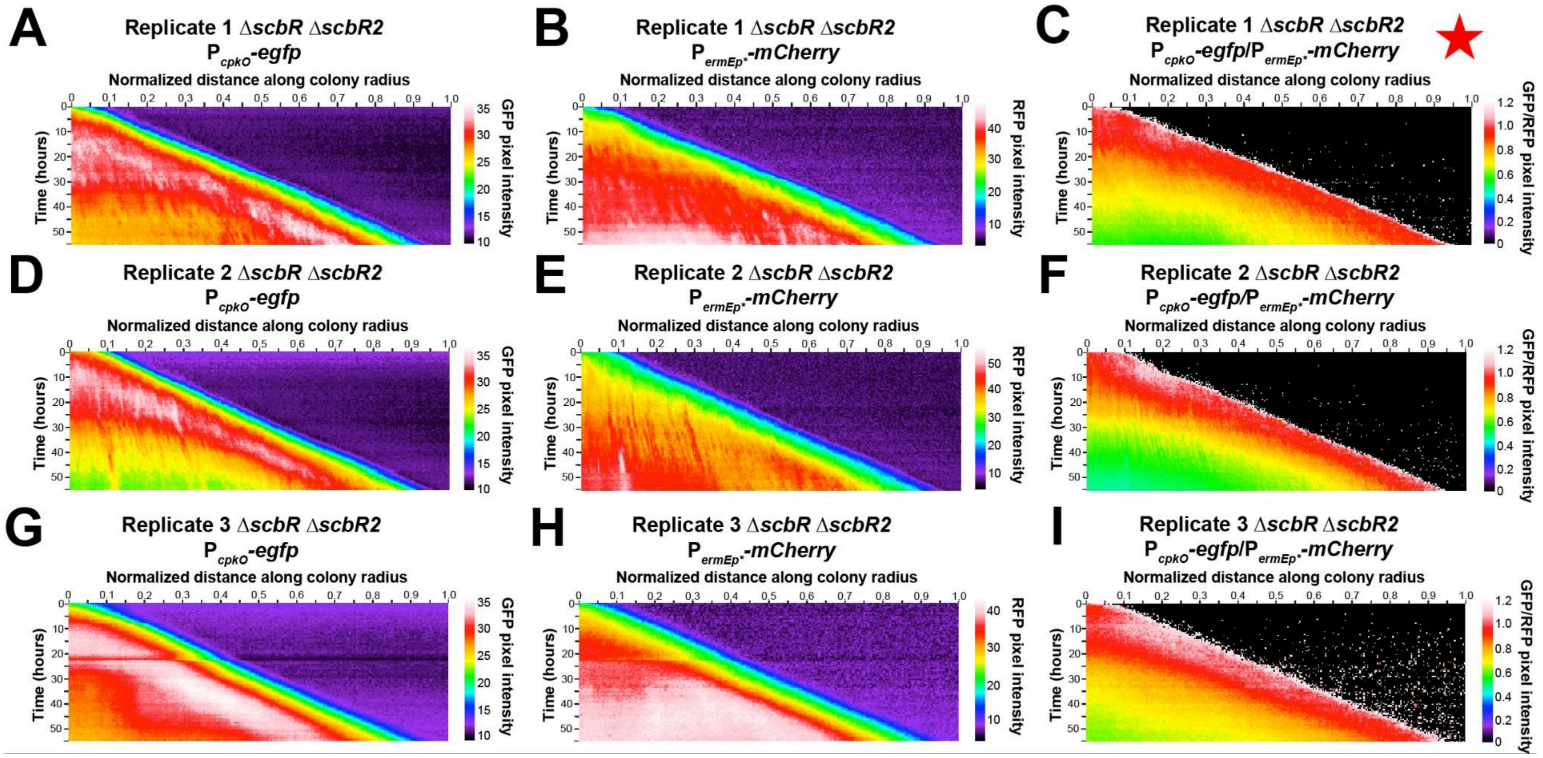


**Figure S14: GFP, RFP, and GFP/RFP kymographs for replicates of Δ*scbR* Δ*scbR2* colonies.** Individual GFP **(A, D, G)**, RFP (**B, E, H)** , and GFP/RFP **(C, F, I)** kymographs for three measured Δ*scbR* Δ*scbR2* colonies (A-C, D-F, G-I). The star indicates the representative kymograph shown in Figure 4F.

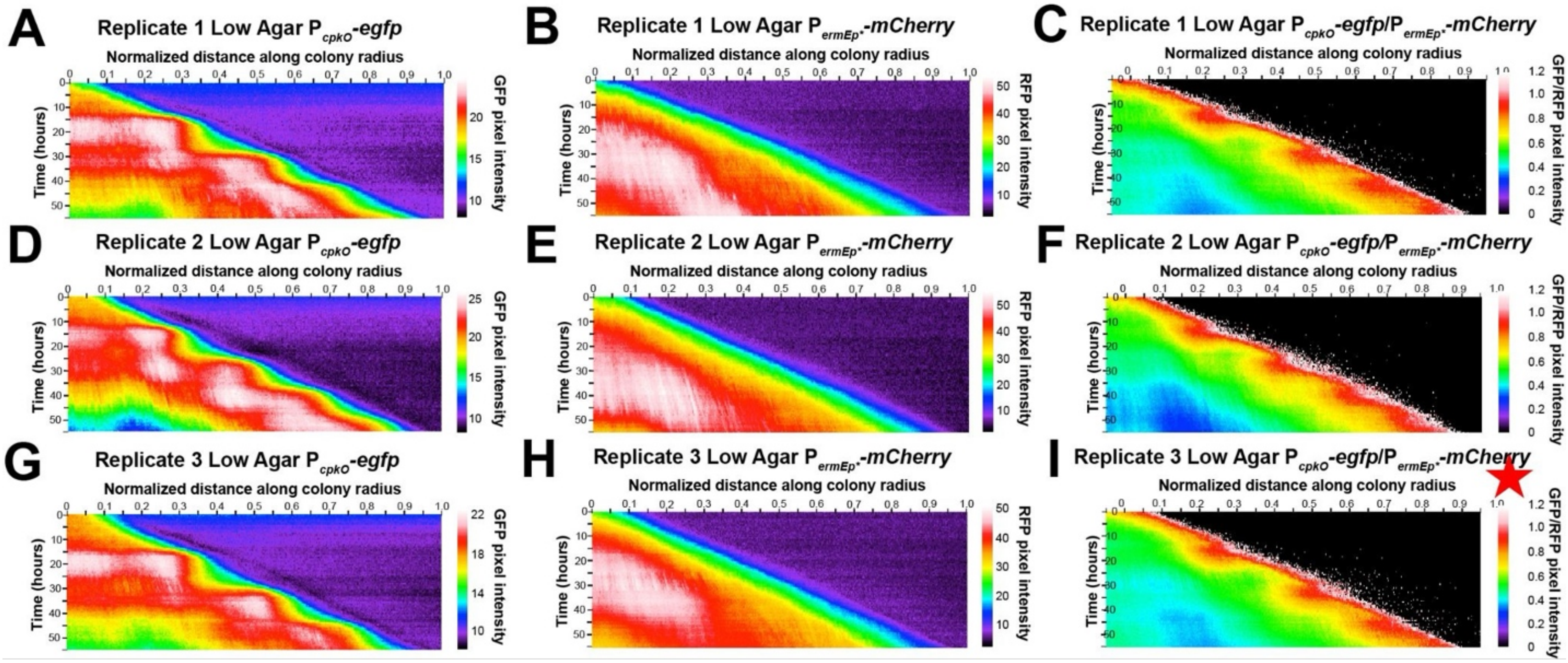


**Figure S15: GFP, RFP, and GFP/RFP kymographs for replicates of WT colonies grown on soft agar.** Individual GFP **(A, D, G)**, RFP **(B, E, H),** and GFP/RFP **(C, F, I)** kymographs for three measured WT colonies grown on soft agar media (A-C, D-F, G-I). The star indicates the representative kymograph shown in Figure 5D.

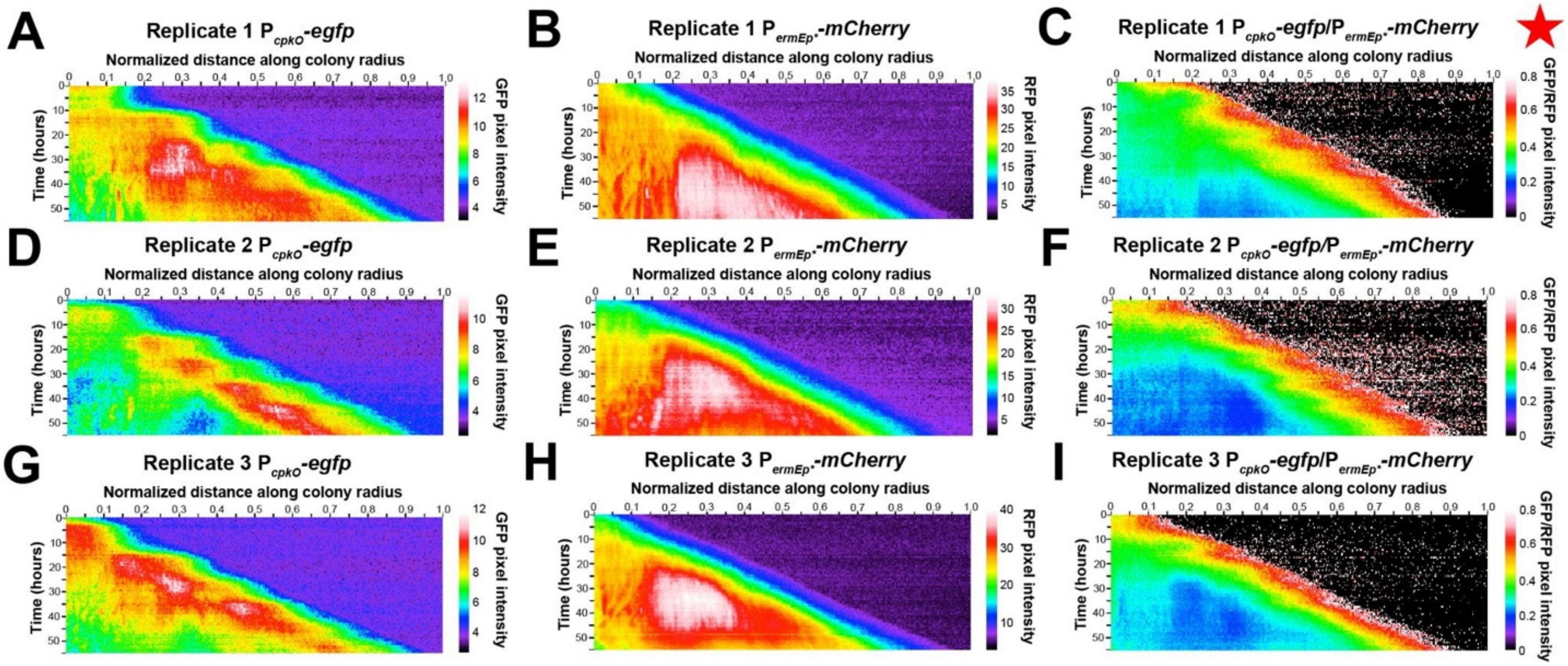


**Figure S16: GFP, RFP, and GFP/RFP kymographs for replicates of WT colonies grown on nutrient-deprived media**. Individual GFP (A, D, G), RFP (B, E, H), and GFP/RFP (C, F, I) kymographs for three measured WT colonies grown on nutrient-deprived media (A-C, D-F, G-I). The star indicates the representative kymograph shown in Figure 5I.

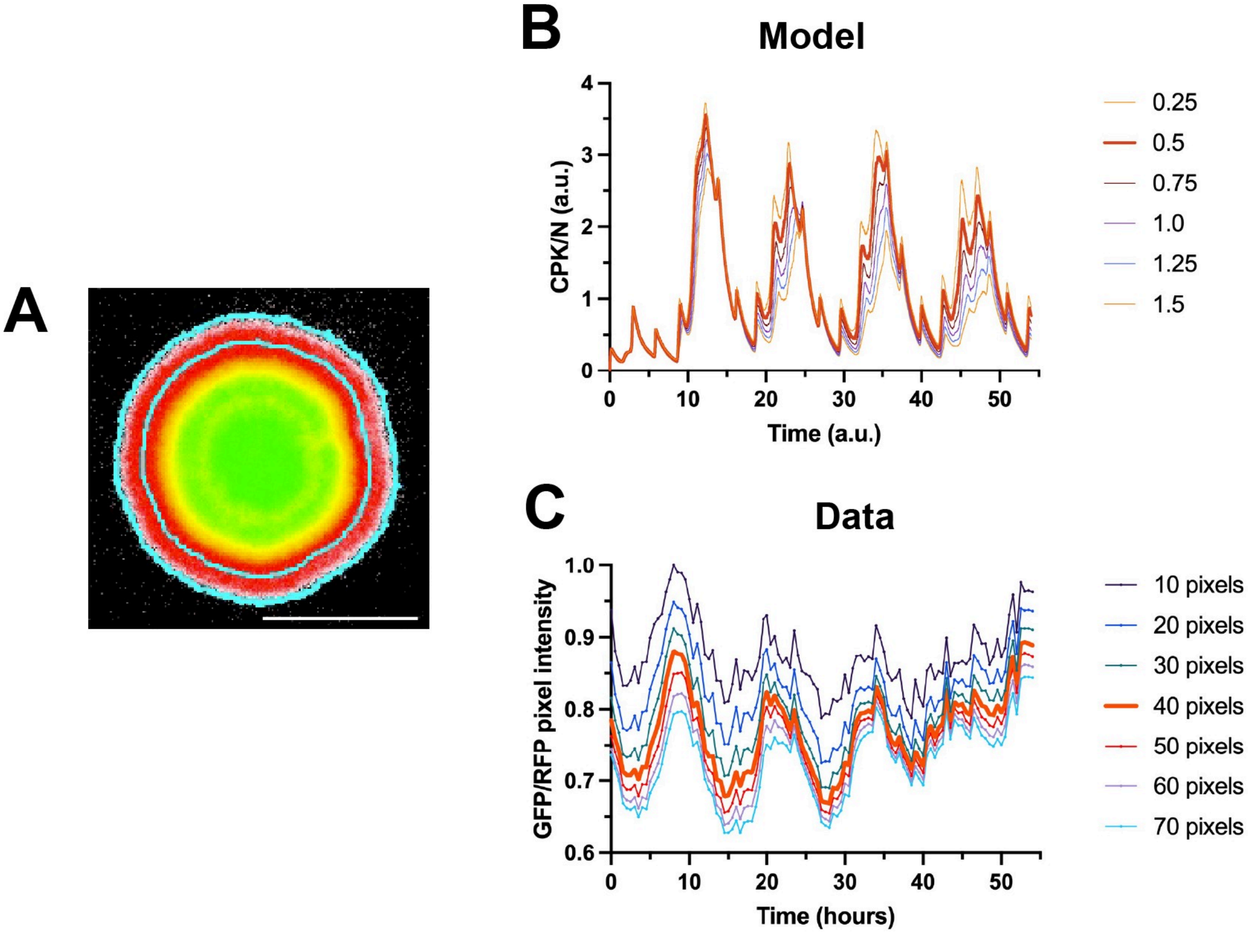


**Figure S17: ROI optimization for plots of normalized *cpkO* expression at the edge of a colony. (A)** Representative image of a WT *S. coelicolor* with an outlined ROI. **(B, C)** A plot charting mean normalized *cpkO* expression in a narrow band at the edge of one simulated (B) or real (C) WT colony. Multiple band sizes were tested and are shown here as individual lines. The band size selected as the standard for this study for simulated and real colonies is bolded in both graphs. (B) Numbers correspond to the value of the ring-width parameter, which controls the size of the band being measured. (C) Numbers correspond to how wide the band measured is, in pixels.

Reaction Diffusion Model of a generic repressilator A $\dashv$ B $\dashv$ C $\dashv$ A circuit:

$$\frac{\partial A}{\partial t} = D_A \nabla^2 A + \alpha_{0,A} + \frac{\alpha_A}{1 + \left(\frac{C}{K_{CA}}\right)^n} - \delta_A\, A$$

$$\frac{\partial B}{\partial t} = D_B \nabla^2 B + \alpha_{0,B} + \frac{\alpha_B}{1 + \left(\frac{A}{K_{AB}}\right)^n} - \delta_B\, B$$

$$\frac{\partial C}{\partial t} = D_C \nabla^2 C + \alpha_{0,C} + \frac{\alpha_C}{1 + \left(\frac{B}{K_{BC}}\right)^n} - \delta_C\, C$$

Reaction Diffusion Model of the *S. coelicolor* QS circuit:

$$\frac{\partial N}{\partial t} = D_N \nabla^2 N + \mu N \left(1 - \frac{N}{K_N}\right)$$

$$\frac{\partial A}{\partial t} = D_A \nabla^2 A + \alpha_A \cdot \frac{N}{1 + \left(\frac{R_2}{K_{R_2 A}}\right)^n} - \delta_A A$$

$$\frac{\partial G}{\partial t} = D_G \nabla^2 G + \alpha_G A - \delta_G G$$

$$\frac{\partial R}{\partial t} = D_R \nabla^2 R + \alpha_R \cdot \frac{N}{1 + \left(\frac{R}{K_{RR}}\right)^n \cdot H(G, \tau)} - \delta_R R$$

$$\frac{\partial R_2}{\partial t} = D_{R_2} \nabla^2 R_2 + \alpha_{R_2} \cdot \frac{N}{1 + \left(\frac{R}{K_{RR2}}\right)^n \cdot H(G, \tau)} - \delta_{R_2} R_2$$

$$\frac{\partial C}{\partial t} = D_C \nabla^2 C + \alpha_C \cdot \frac{N}{1 + \left(\frac{R_2}{K_{R_2 C}}\right)^n + \left(\frac{R}{K_{RC}}\right)^n \cdot H(G, \tau)} - \delta_C C$$

**Figure S18: A mathematical model of a standard repressilator circuit. (Top)** A set of PDEs that simulate a standard repressilator circuit that incorporates spatial diffusion. **(Bottom)** The mathematical model developed for this study that simulates the *S. coelicolor* QS circuit.

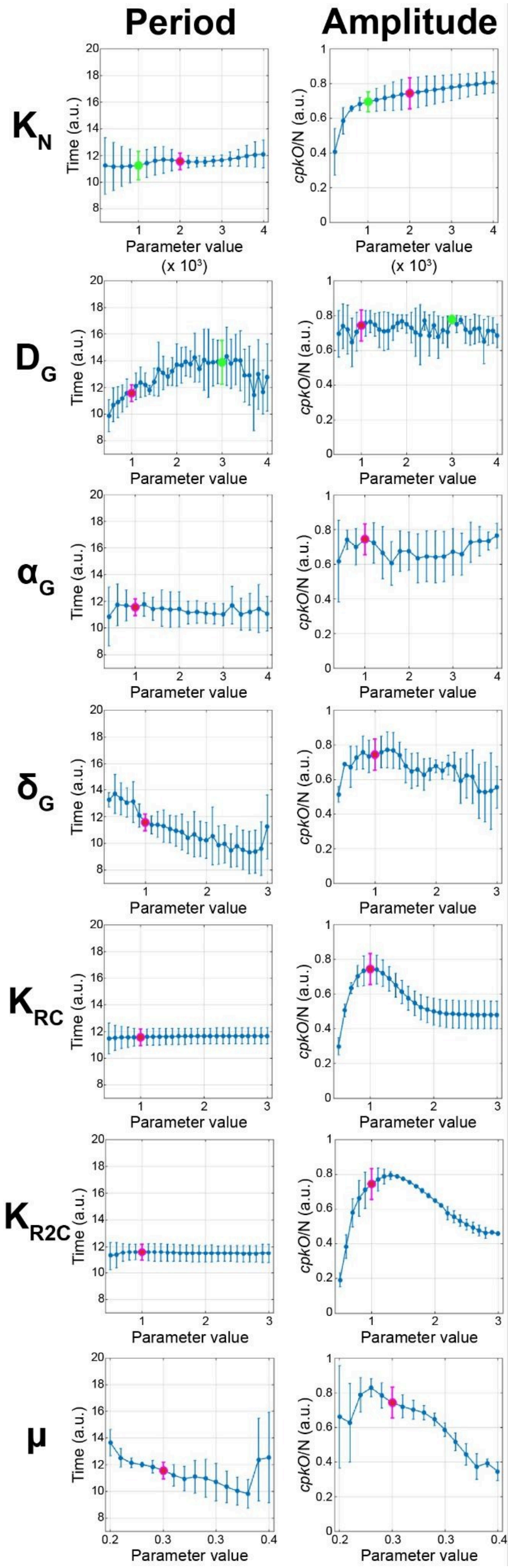
Period
Amplitude
K_N
D_G
α_G
δ_G
K_RC
K_R2C
μ
Time (a.u.)
cpkO/N (a.u.)
Parameter value
(x 10³)

**Figure S19: Sensitivity analysis on the mathematical model.** Measured average periodicity **(left)** and amplitude **(right)** of simulated normalized *cpkO* expression pulses at the virtual colony edge at several tested parameter values of $K_N$, $D_G$, $\alpha_G$, $\delta_G$, $K_{RC}$, $K_{R2C}$, and μ. Data points are an average measurement of all oscillations observed during the corresponding simulation, with error bars representing standard deviation. Measurements were taken on datasets that were filtered to obtain clear pulses (see the Materials and Methods for details regarding this filtering). Points highlighted in magenta indicate the nominal value of each parameter. Points highlighted in green indicate the value to which the parameter was adjusted for further *in silico* and *in vitro* analysis.

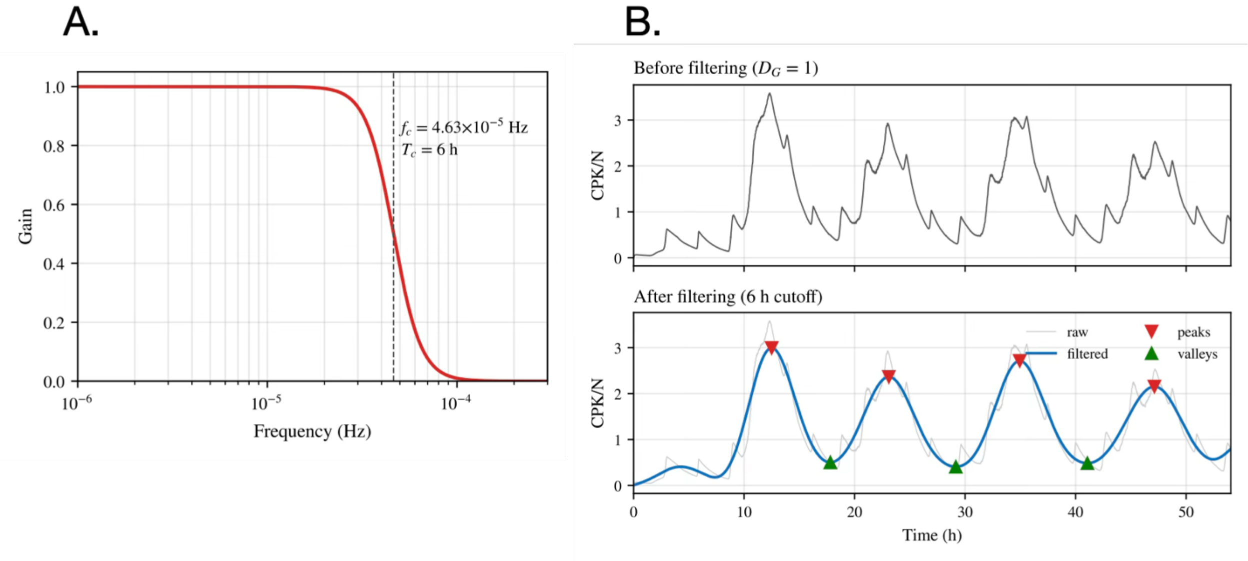


**Figure S20: Filtering *in silico* data to resolve high-frequency oscillations. (A)** The third-order butterworth filter applied to the model output. The dashed line indicates the cutoff value. **(B, top)** Original normalized simulated *cpkO* expression at the colony edge. **(B, bottom)** The same dataset, with the filter applied. Red triangles indicate the peaks as identified automatically by the MATLAB Signal Processing Toolbox. Green triangles indicate the valleys, identified by the same software. Amplitude and periodicity as assessed in the sensitivity analysis were measured on datasets with this applied filter.

| Parameter | Value | Description |
| --- | --- | --- |
| n | 10 | Sharpness of repression |
| $\tau$ | 0.1 | QS threshold |
| $\mu$ | 0.3 | Growth rate |
| k | 100 | Sharpness of QS filter |
| *$K_N$* | *2000* | *Carrying capacity* |
| $D_N$ | 10000 | Colony spread over space |
| eps | 0.01 | Sharpness of the colony edge |
| *$D_G$* | *1* | *Diffusion of GBLs* |
| $\alpha_A$ | 1 | Expression of *scbA* |
| **$\alpha_G$** | **1** | **Production of GBLs** |
| **$\alpha_R$** | **1.1** | **Expression of *scbR*** |
| **$\alpha_{R2}$** | **1.8** | **Expression of *scbR2*** |
| **$\alpha_C$** | **3** | **Expression of *cpkO*** |
| $\Delta_A$ | 1 | Degradation of *scbA* |
| **$\Delta_G$** | **1** | **Degradation of GBLs** |
| $\Delta_R$ | 0.1 | Degradation of *scbR* |
| $\Delta_{R2}$ | 0.5 | Degradation of *scbR2* |
| **$\Delta_C$** | **0.25** | **Degradation of *cpkO*** |
| $K_{R2A}$ | 0.4 | Half-saturation constant for ScbR2 on *scbA* |
| $K_{RR}$ | 0.5 | Half-saturation constant for ScbR on *scbR* |
| $K_{RC}$ | 1 | Half-saturation constant for ScbR on *cpkO* |

| | | |
|---|---|---|
| $K_{RR2}$ | 0.5 | Half-saturation constant for ScbR on *scbR2* |
| $K_{R2C}$ | 1 | Half-saturation constant for ScbR2 on *cpkO* |

**Table 1. List of parameters incorporated into the model.** Bolded parameters represent those that were optimized using experimental data generated in this study. Underlined parameters represent those that were adjusted based on biological truths. Italicized parameters represent those that were altered experimentally in Figure 6.

| Strain | Relevant information | Origin/reference |
|---|---|---|
| <u>*S. coelicolor* strains</u> | | |
| M145 $P_{cpkO}$-*egfp*, $P_{ermEp*}$-*mCherry* | Wild-type strain, *cpkO* reporter | (73) |
| M145 Δ*scbA* $P_{cpkO}$-*egfp*, $P_{ermEp*}$-*mCherry* | M145 derivative, markerless *scbA* deletion, *cpkO* reporter | This study |
| M145 Δ*scbR* $P_{cpkO}$-*egfp*, $P_{ermEp*}$-*mCherry* | M145 derivative, markerless *scbR* deletion, *cpkO* reporter | This study |
| M145 Δ*scbR2* $P_{cpkO}$-*egfp*, $P_{ermEp*}$-*mCherry* | M145 derivative, markerless *scbR2* deletion, *cpkO* reporter | This study |
| M145 $P_{bldN}$-*egfp*, $P_{ermEp*}$-*mCherry* | Wild-type strain, *bldN* reporter | This study |
| M145 $P_{scbA}$-*egfp*, $P_{ermEp*}$-*mCherry* | Wild-type strain, *scbA* reporter | This study |
| <u>*E. coli* strains</u> | | |
| XL1-BLU | *recA1 endA1 gyrA96 thi-1 hsdR17 supE44 relA1 lac [F´ proAB lacIqZΔM15 Tn10 (Tetr)]* | Agilent Technologies |
| ET12567/pUZ8002 | F- *dam*-13::Tn9 *dcm*-6 *hsdM hsdR zjj*-202::Tn10 *recF*143 *galK2 galT22 ara*-14 *lacY*1 *xyl*-5 *leuB6 thi*-1 *tonA*31 *rpsL*136 *his*G4 *tsx*-78 *mtl*-1 *gln* | (61, 62) |

**Table 2. List of strains used in this study**

**Movie S1 (separate file): Fluorescence time-lapse movie of $P_{cpkO}$-*egfp*/$P_{ermEp*}$-*mCherry* (GFP/RFP) expression in a representative WT colony.** The time-lapse movie spans 54 hours, starting from a colony 42 hours old. Each frame of the movie is an image captured every 30 minutes.

**Movie S2 (separate file): Simulated *cpkO*/N in a virtual colony.**
